# Partitioning of ribonucleoprotein complexes from the cellular actin cortex

**DOI:** 10.1101/2021.10.01.462753

**Authors:** Isaac Angert, Siddarth Reddy Karuka, Louis M. Mansky, Joachim D. Mueller

## Abstract

The cell cortex plays a crucial role in cell mechanics, signaling, and development. However, little is known about the influence of the cortical meshwork on the spatial distribution of cytoplasmic biomolecules. Here, we describe a new fluorescence microscopy method to infer the intracellular distribution of labeled biomolecules with sub-resolution accuracy. Unexpectedly, we find that RNA-binding proteins are partially excluded from the cytoplasmic volume adjacent to the plasma membrane that corresponds to the actin cortex. Complementary diffusion measurements of RNA-protein complexes suggest that a rudimentary model based on excluded volume interactions can explain this partitioning effect. Our results suggest the actin cortex meshwork may play a role in regulating the biomolecular content of the volume immediately adjacent to the plasma membrane.

**Teaser:** A novel microscopy method reveals exclusion of RNA-protein complexes from the actin cortex due to their large hydrodynamic size.

## Introduction

Both the actin cortex and RNA are fundamental building blocks of eukaryotic cells. The cortex, acting in concert with other cytoskeletal elements, is involved in nearly all aspects of cell mechanics, such as cell structure, membrane rigidity, migration and division (*1*). Due to its location adjacent to the plasma membrane, the cortex plays important roles in signal transduction and mechanical response pathways (*1, 2*). In contrast, RNA molecules serve as genomic messengers and constitute major components of the cellular translational machinery. Specific localization of mRNA to sub-cellular regions has been widely observed (*3*). RNA localization to sub-cellular compartments such as the endoplasmic reticulum (ER), processing (P)-bodies or distal cytoplasm in neurons is critical to aspects of development (*4*) and is an important regulator of cellular function (*5*). Due to their highly structured and extended nature, mRNAs have hydrodynamic radii that are dramatically larger than those of the globular proteins they encode (*6*). Based on a recent study (*7*), a typical mRNA of 1 – 2 kilobase length (*8, 9*) is expected to have a hydrodynamic radius in the range of ∼10 nm. The radii of ribosomes are of comparable size (10 – 15 nm) (*10, 11*).

Previous biophysical studies have established that the cell cytoplasm is a complex medium (*12, 13*) that modulates the diffusion and localization of large (>10 nm) particles in a size-dependent manner (*14*–*18*). These observations have prompted the question of whether the large size of the cellular translational machinery might broadly influence its subcellular localization (*19*). Here, the novel observation that large riboneucleoprotein (RNP) complexes are partially excluded, or partitioned, from the interior of the actin cortex is reported. These observations are facilitated by recent developments in two-photon dual-color (DC) z-scan fluorescence microscopy (*20*), which is adapted from its previous application quantifying binding of peripheral membrane proteins to now infer protein localizations inside living cells with sub-diffraction resolution of lengths. DC z-scans are acquired by moving the two-photon excitation volume vertically through the cytoplasm of adherent cells. This movement of the excitation volume yields fluorescent intensity traces that encode the vertical intracellular distribution of a fluorescently labeled protein species as well as that of a distinctly colored reference fluorescent protein. Although different reference species are possible, our typical choice is a soluble, purely cytoplasmic protein, which serves as a “paint” that marks the cytoplasmic extent. Differential vertical localization between the fluorescently labeled protein and the reference protein is encoded as subtle differences in the shape of the DC z-scan traces. The magnitude of the differential vertical localization is quantitatively recovered from the data by fitting the z-scan traces. In a typical sample, this procedure attains a resolution of 20 – 50 nm in after collection of 60 s of z-scan data.

## Results

### DC z-scan measures sub-diffraction distances in living cells

Initial proof-of-principle experiments were conducted with U2OS cells to demonstrate the utility of DC z-scan to quantify sub-diffraction differences in intracellular protein localizations. For detailed descriptions of the z-scan model geometries used in the following, see the Materials and Methods section. First, cells were co-transfected with mCherry-RXR (a protein that localizes in the nucleus) and EGFP (which diffuses freely throughout the cell). DC z-scans through the cell nucleus were fit to the S^G^-S^CH^ (Eq. 13) model to identify the slab length associated with the mCherry fluorescence, which corresponds to the nuclear thickness, and the slab length of the EGFP signal, which corresponds to the slightly longer thickness of the entire cell (Fig. 1A). A single representative z-scan, acquired in 5 s and subsequently fitted, shows that the mCherry-RXR and EGFP z-scans are well described by this model, with flat residuals and χ_v_^2^ = 1.2 (Fig. 1B). The fit identified the intensity amplitudes of mCherry-RXR and EGFP, the locations of the bottom and top cell edges (*a*^*G*^ = 2.16 µm and *b*^*G*^ = 5.80 µm), as well as the locations of the bottom and top nuclear edges (*a*^*CH*^ = 2.51 µm and *b*^*CH*^ = 5.03 µm), respectively. The distances between the edges of the EGFP and mCherry slab were defined at the top and bottom of the cell by *Δ*^*T*^ = *b*^*CH*^ - *b*^*G*^ = -770 nm and *Δ*^*B*^ = *a*^*G*^ - *a*^*CH*^ = -350 nm, respectively. Since *Δ* measures the displacement between the borders of the EGFP and mCherry layers, its value can be positive or negative in general. Distances are reported by taking the absolute value, which in this instance determines the thickness of the cytoplasmic layers surrounding the top and bottom of the nucleus (Fig. 1A). To improve the resolution and identify statistical uncertainties, 60 s of repeated z-scans were then acquired at the same location within this cell and subsequently fit to obtain an average of *Δ*^B^ = -410 ± 16 nm and *Δ*^T^ = -800 ± 15 nm (SEM from 12 scans). Multiple 60 s DC z-scan measurements were then acquired in multiple cells. Each of these experiments yielded estimates of *Δ*^B^ and *Δ*^T^ at each location within a cell. These values are summarized by a histogram (Fig. 1C) and the resulting distribution is characterized by the averages (see Materials and Methods) <*Δ*^*T*^> = -340 ± 50 nm and <*Δ*^*B*^*>* = -260 ± 30 nm (SEM, *n* = 19, where *n* is the number of scan locations averaged).

**Fig. 1:**
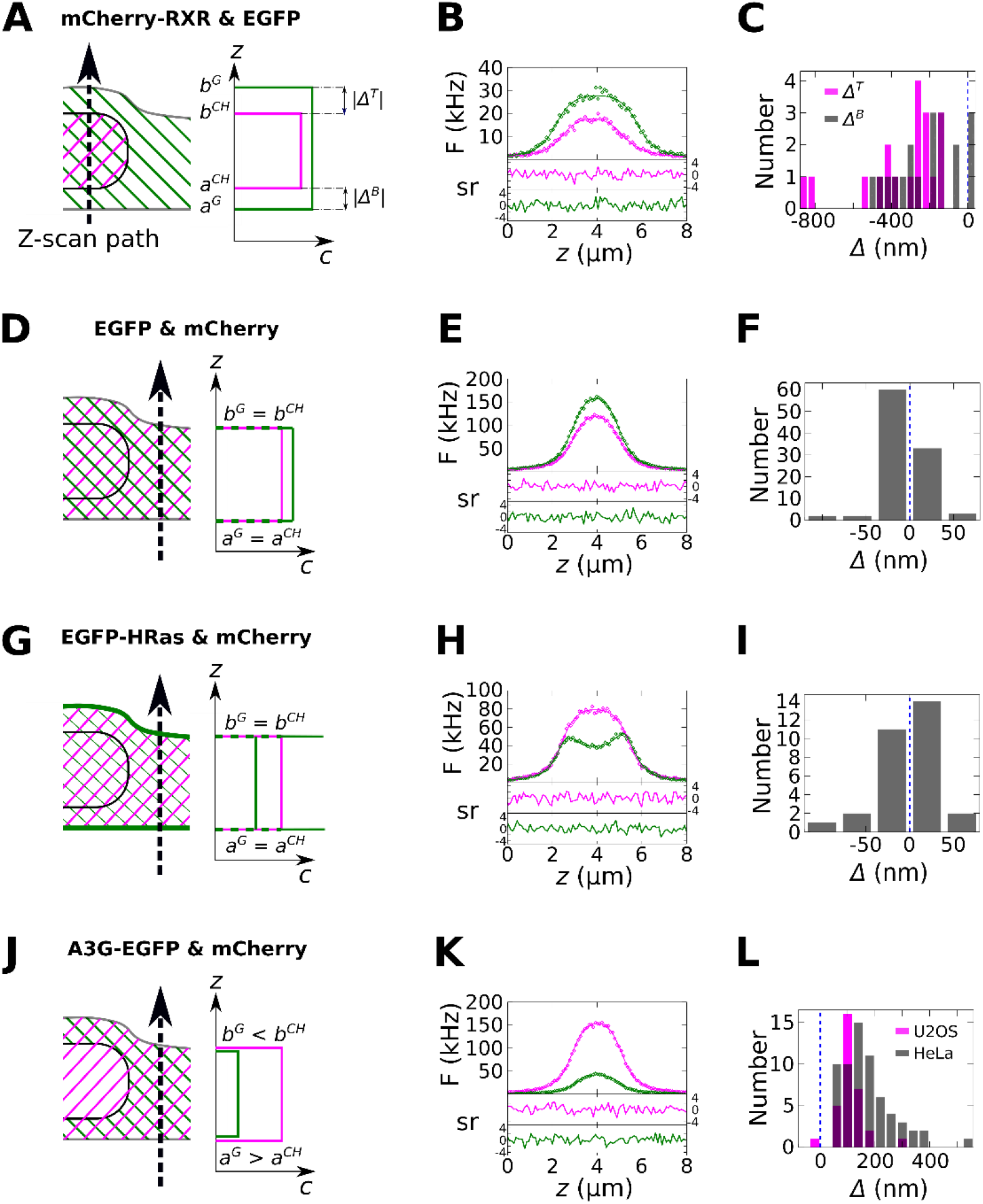
DC z-scans in living cells. The z-scan trajectory through the cell is depicted in the cartoons along with crosshatched regions indicating the subcellular localization of the EGFP- and mCherry-labeled proteins (A, D, G, J, left panel) together with the concentration profile indicated by z-scan fitting for the EGFP (green lines) and mCherry (magenta lines) protein species (A, D, G, J, right panel). B, E, H, K) Fits (solid lines) of representative z-scan intensity traces of the green (green diamonds) and red (magenta diamonds) detection channel are shown along with their normalized residuals *sr*. The cell edge locations *a*^*G*^, *b*^*G*^, *a*^*CH*^, *b*^*CH*^ extracted from the fits determine the vertical distance *Δ* separating the EGFP- and mCherry-labeled slab edges in each z-scan. C, F, I, L) Histograms of the average *Δ* values estimated from each scan location in many cells. Experiments with different EGFP- and mCherry-labeled protein species are indicated in rows: (A, B, C) EGFP and mCherry-RXR, (D, E, F) EGFP and mCherry, (G, H, I) EGFP-HRas and mCherry, and (J, K, L) A3G-EGFP and mCherry.

Importantly, the DC z-scan experiments reported in the present work were performed using two-photon microscopy, which allows EGFP and mCherry to be co-excited by using the same laser focus. This results in red and green channel point spread functions (PSFs) that are identical at the nanometer scale (Fig. S1). This lack of chromatic shift can be seen in control DC z-scans that were performed through the cytoplasm of U2OS cells co-expressing EGFP and mCherry fluorescent proteins (Fig. 1D). Since both proteins are soluble species within the cell cytoplasm, DC z-scans are expected to recover *Δ =* 0, on average. As shown in Fig. 1E, a representative DC z-scan fits to a S^G^-S^CH^ (Eq. 13) model with flat residuals and χ_v_^2^ = 1.1. Repeated DC z-scans in multiple cells recovered mean values of <*Δ*^*B*^> = -2.3 ± 1.2 nm and <*Δ*^*T*^*>* = 1.0 ± 3.1 nm (SEM, *n* = 59). In the absence of a statistically significant difference between the results from the bottom and top of the cell, the data were combined, and the superscript dropped. The histogram of the combined displacements (Fig. 1F) determined <*Δ*> = -0.5 ± 1.7 nm (SEM, *n* = 118).

Next, DC z-scans were performed in U2OS cells co-expressing EGFP-HRas and mCherry (Fig. 1G). EGFP-HRas was previously reported to be a peripheral membrane protein that binds to the plasma membrane as a monomer (*21*). Previous z-scan studies established that the axial distribution of EGFP-HRas in the cytoplasm and at the PM was successfully modeled by the δSδ layer geometry (*21, 22*), where the two δ layers account for PM bound protein and the S layer accounts for protein in the cytoplasm. Thus, DC z-scans from cells co-expressing mCherry and EGFP-HRas were modeled by the (S^CH^-δSδ^G^) geometry (Eq. 13). A fit of a representative z-scan attained good quality of fit with χ_v_^2^ = 1.2 and flat residuals. Repeated z-scans in the cytoplasm of multiple cells co-expressing EGFP-HRas and mCherry (Fig. 1I) determined a mean displacement of <*Δ>* = -2.6 ± 5.4 nm (SEM, *n* = 30), which is consistent with the expected value of *Δ* = 0. This result indicated that the location of the cytoplasm slab edge, as determined by fitting the soluble mCherry species to the S model, coincided with the location of the PM, as determined by fitting the EGFP-HRas species to the δSδ model.

### APOBEC 3G is depleted from a thin volume adjacent to the PM

Next, the axial cytoplasmic distribution of APOBEC 3G (A3G) labeled with EGFP and co-expressed with mCherry (Fig. 1J) was examined. A3G is part of the innate immune system and acts as a cytidine deaminase that restricts replication of Vif-negative HIV-1. A3G binds strongly to both cellular and viral RNAs, the latter of which is essential to its retroviral restriction activity (*23, 24*). A representative z-scan acquired in the cytoplasm of a HeLa cell showed a single broad peak for both mCherry and A3G-EGFP (Fig. 1K). Since the trace shape was suggestive of a slab geometry, both species were fit to slab models (S^CH^-S^G^, Eq. 13). This fit provided a good description of the DC z-scan intensity profiles with *χ*_*v*_^*2*^ = 1.3 and flat residuals (Fig. 1K). Interestingly, this fit identified non-zero values for the displacements at the top and bottom PM (*Δ*^*B*^ = 184 nm and *Δ*^*T*^ = 167 nm). Repeated z-scan acquisitions at the same location in this cell determined average values *Δ*^B^ = 180 ± 6 nm and *Δ*^T^ = 162 ± 12 nm (SEM from 12 scans). This surprising observation suggests that the thickness of the soluble A3G-EGFP layer is less than the thickness of the mCherry layer. Repeated z-scan acquisitions in multiple HeLa cells (Fig. 1L) found an average of <*Δ*>_HeLa_ = 155 ± 10 nm (SEM, *n* = 64), indicating that A3G does not occupy the entire cytoplasmic volume. To reflect these results *Δ* is also referred to as the effective depletion length. To further validate this observation, DC z-scan measurements were collected using a different cell line (U2OS), which resulted in an average of <*Δ*>_U2OS_ = 110 ± 9 nm (SEM, *n* = 32). Rescaled histograms of Fig. 1 are shown in Fig. S2 to facilitate visual comparison of the different samples. Because the uncertainty in the measured depletion length may arise from intracellular or intercellular variability, the variation of the depletion length at distinct locations within the same cell and between multiple cells was determined (Table S1), which indicates that the majority of the variability is accounted for by intracellular heterogeneity. Additionally, the data in Table S1 demonstrate that DC z-scan attains a resolution of 20-50 nm in typical samples after acquisition of 60 s of z-scan data.

### PM adjacent depletion is reduced by disrupting F-actin

Intriguingly, this result implies that cytoplasmic A3G-EGFP was excluded from regions adjacent to the PM with apparent thickness of 100 - 200 nm. Given the magnitude of this value, further experiments were conducted to test whether A3G-EGFP might be partially excluded from the dense meshwork of the actin cortex. To do this, HeLa cells were treated with ethanol, which disrupts actin assemblies and leads to the formation of large plasma membrane blebs (*25*) (Fig. 2A) that lack an intact actin cortex (*26, 27*). A representative z-scan performed near the center of these blebs showed good quality of fit to a S^CH^-S^G^ model (Eq. 13 and Fig. 2C). Repeated z-scans in blebs from HeLa cells co-expressing A3G-EGFP and mCherry (Fig. 2D) determined an average of <*Δ*> = 31 ± 10 nm (SEM, *n* = 14), which differs from the results obtained in untreated HeLa cells by 8.8 SD.

**Fig. 2:**
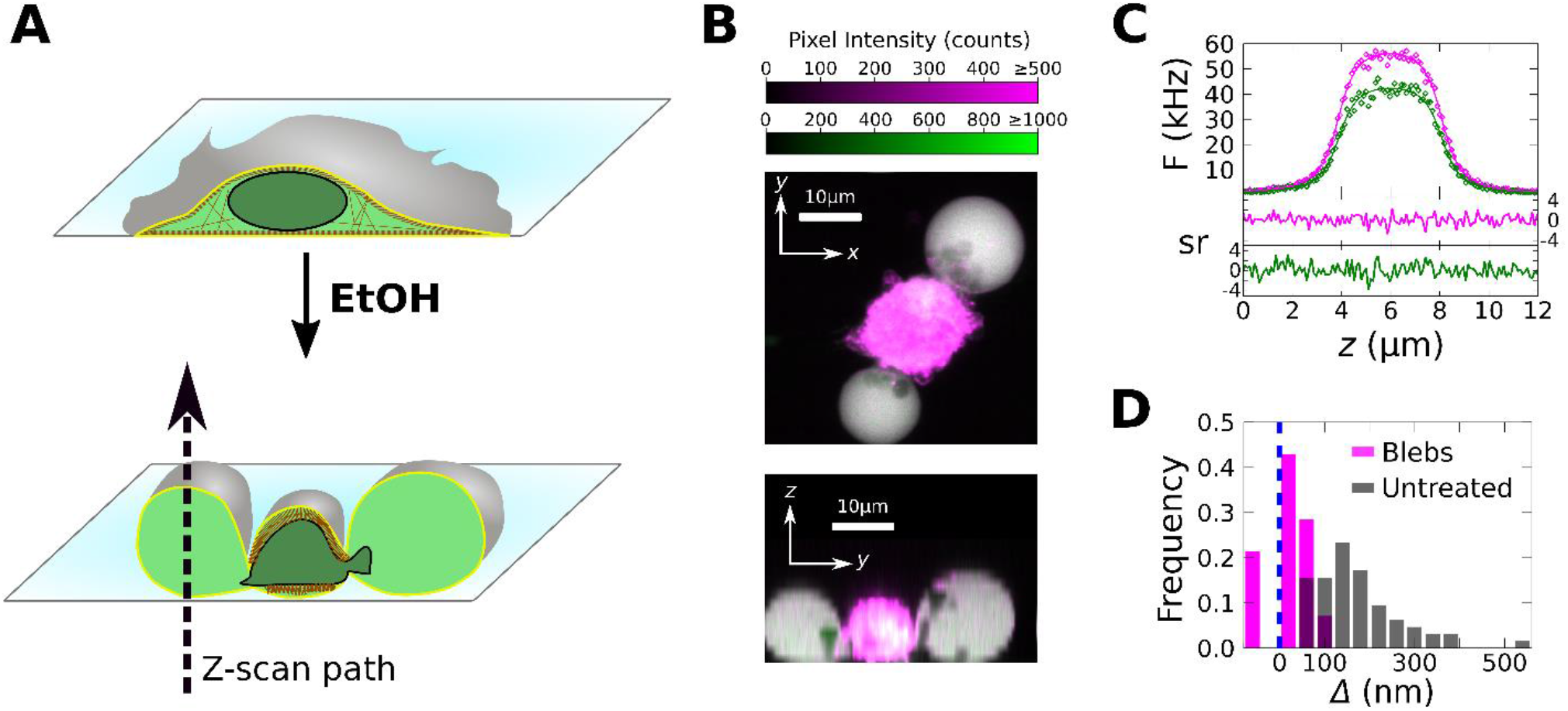
Z-scans in blebs produced from HeLa cells co-expressing A3G-EGFP and mCherry. A) A graphic depicting the production of large PM blebs from adherent cells via ethanol treatment is shown. In this example, the cytoskeleton is denoted by brown colors. B) A DC 2-photon laser scanning z-stack image of a HeLa cell with two large PM blebs is shown. The x-y image was created by the summation of the z-stack over the z-dimension. The y-z image was produced by the sum of the x-dimension over two volume-sections passing roughly through the center of each bleb. While significant red autofluorescence is produced by the ethanol treatment near the cell center, it is typically absent in the blebs. Z-scan locations were selected to pass near the center of each bleb, while avoiding regions possessing autofluorescence or identifiable structures. C) Fit of a representative z-scan passing through a HeLa cell bleb (*χ*_*v*_^*2*^= 1.2). D) Histogram of the average *Δ* values estimated from each scan location in the cytoplasm of many HeLa cells not treated with ethanol (magenta bars) and in blebs produced by ethanol treatment (gray bars).

Because the ethanol treatment may induce pleiotropic effects, it was verified that blebs induced in HeLa cells expressing A3G-EGFP by cytochalasin D treatment showed a similar reduction in the depletion effect (Fig. S3). To provide additional support for the role of the cortex in PM adjacent depletion, DC z-scans were performed in HeLa cells treated with latrunculin A (LA), a toxin which prevents actin polymerization. In contrast to experiments with cytochalasin D, LA was used at a concentration low enough to maintain gross cell morphology (i.e., below the bleb-forming concentration). DC z-scans performed in the cytoplasm of LA treated cells indicated a reduction in <*Δ*> compared to a solvent control (Fig. S3). Taken together, the strong reduction in *Δ* found in blebs and the decrease observed in the presence of latrunculin A support the conclusion that PM adjacent depletion is driven by the presence of the actin cortex.

### The depletion length depends on the cortical thickness

Given that dependence of cortical architecture on the cell type has been observed (*28*–*30*), the differences in effective depletion length observed between HeLa and U2OS cells might reflect natural differences in cortex thickness between these cell types. To investigate this, direct measurements of the cortex thickness were performed using distinct fluorescent colors to mark the actin cortex and the plasma membrane. A previous study used this approach to estimate the actin cortex thickness from the displacement between intensity peaks corresponding to the actin and the PM marker in confocal images (*31*). In the present study, two-photon DC z-scan was employed instead of confocal imaging to determine the cortex thickness from the intensity profiles of EGFP-HRas (which binds to the PM) and Lifeact-mApple (which binds to actin and marks the cortex) (*32*) by fitting to on a quantitative model of the entire intensity traces (Fig. 3A). The axial concentration profile of both labels is depicted in Fig. 3B, which shows an enrichment of EGFP-HRas and Lifeact-mApple at the PM and within the cortex, respectively. A small amount of both fluorescent proteins is also present in the cytoplasm. Based on this profile, EGFP-HRas is again modeled by the δSδ geometry. The distance separating the two HRas δ layers represents the cell thickness *L*_cell_.

**Fig. 3:**
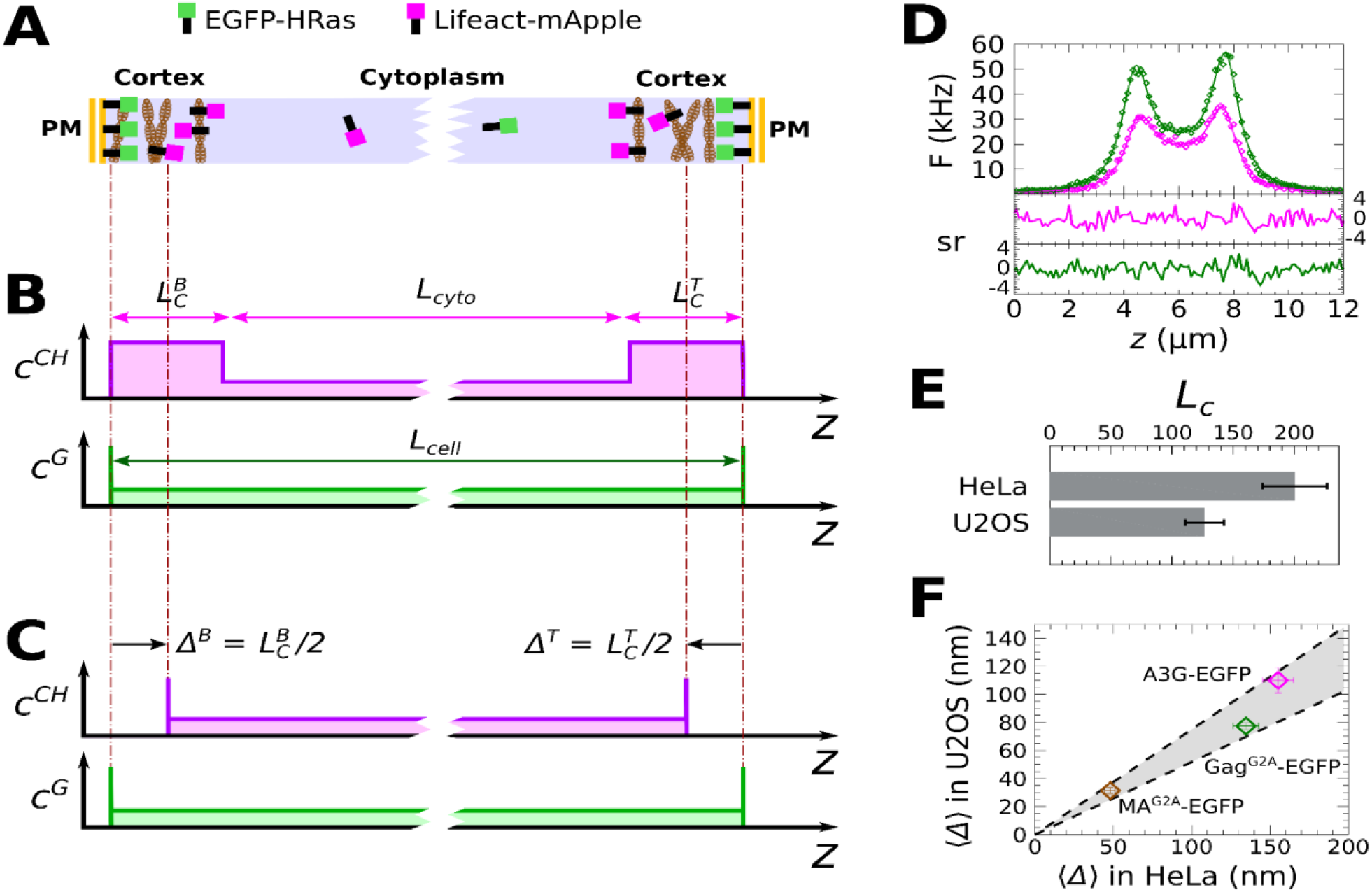
DC z-scan measurement of cortex thickness. A) Lifeact-mApple demarcates the actin cortex, and EGFP-HRas demarcates the PM location. These fluorescent proteins lead to the SSS^CH^-δSδ^G^ concentration profile depicted in (B). C) DC z-scans were fit to a δSδ^CH^-δSδ^G^ model. The location of the mApple δ layers corresponds to the centroid of the cortical slabs, so that the effective depletion length is half the thickness of the cortex slab. D) A fit of a DC z-scan through a HeLa cell co-expressing EGFP-HRas and Lifeact-mApple. E) Average cortex thickness values estimated by DC z-scan in HeLa and U2OS cells. F) <*Δ*> is plotted in U2OS cells versus the same vales in HeLa cells for MA^G2A^-EGFP (brown), Gag^G2A^-EGFP (green) and A3G-EGFP (magenta). The ratio of cortex thicknesses in the cell lines corresponds to the slope of the data as is indicated by the shaded area and dashed lines, which correspond to the slope-uncertainty derived from the data in panel (E).

The concentration profile of Lifeact-mApple implies a wide cytoplasmic slab of length *L*_*cyto*_ separating two thin slabs representing the cortex at the bottom and the top membrane with thickness 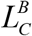 and 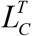 (Fig. 3B), giving rise to an SSS^CH^ (triple S-layer) geometry. A uniform slab with a sharp transition between the cortical and cytoplasmic spaces model of the cortex has been previously used (*31*). Because the expected slab length *L*_*C*_ of the cortex is smaller than 500 nm, its length is experimentally indistinguishable from a δ layer (*33*) by single-color (SC) z-scans. However, the centroid position of the cortical layer is expected to coincide with the midpoint of the slab as depicted in Fig. 3B. Given this, modeling the Lifeact-mApple species as a δSδ^CH^ geometry can identify the cortex centroid from the fitted location of the δ layers. Combined with the EGFP-HRas species, this DC z-scan fit (δSδ^CH^-δSδ^G^) identifies the locations of the δ layers corresponding to the cortex centroid and the PM as well as their displacement *Δ*. The distance |*Δ*| separating both δ layers is half of the actin cortex thickness, |*Δ*| *= L*_*C*_ / 2 (Fig. 3B). This relation was verified by fitting simulated DC z-scan intensity profiles from the SSS^CH^-δSδ^G^ geometry to the δSδ^CH^-δSδ^G^ model (Fig. S4). Repeated DC z-scans were then collected in cells co-expressing EGFP-HRas and Lifeact-mApple proteins in order to determine cortical thickness. A representative DC z-scan and fit to the δSδ^CH^- δSδ^G^ model (Fig. 3C) for both proteins is shown in Fig. 3D. Data taken from multiple cells resulted in cortex thickness estimates of 126 ± 16 nm (SEM, *n* = 7) for U2OS cells and 200 ± 24 nm (SEM, *n* = 15). for HeLa cells (Fig. 3E).

Since both the depletion length *Δ* for A3G-EGFP and the cortex thickness differed between the two cell lines, *<Δ>* in HeLa and U2OS cells were plotted for comparison (Fig. 3F). Additional DC z-scans were performed in cells expressing HIV-1 Gag^G2A^-EGFP and its matrix (MA) domain, MA^G2A^-EGFP, with mCherry again as a reference marker of the cytoplasmic volume for each instance. These *<Δ>* values were added to Fig. 3F (for histograms, see Fig. S5). Interestingly, all 3 data points cluster near a line of slope 0.66, indicating that, although the magnitude of the depletion effect varies for each protein, it is systematically scaled by ∼66% between these two cell lines. This same ratio corresponds well to the cortex thickness ratio estimated by DC z-scan between HeLa and U2OS cells (0.66 ± 0.11, shaded area in Fig. 3F), suggesting that the cell-type dependent differences in *<Δ>* are due to differences in their cortical actin thicknesses.

### PM adjacent depletion of proteins is driven by RNA-binding

All 3 proteins in Fig. 3F are known to be RNA-interacting proteins: HIV-1 Gag^G2A^ binds specifically and non-specifically to RNA sequences through basic residues and two zinc fingers in its nucleocapsid (NC) domain (*34*–*36*), while the MA domain of Gag separately interacts with RNA via a stretch of basic residues in its highly basic region (HBR) (*37*) and has preferential tRNA binding affinity (*38*). A3G interacts with RNA primarily through its N-terminal domain zinc finger (*23, 24*). To test whether the depletion effect is specific to RNA-binding proteins, DC z-scans were conducted with the GABA-A receptor associated protein fused to EGFP (GABARAP-EGFP) and a tandem hexamer of the Venus fluorescent protein, Venus_6_. Both proteins lack a known RNA binding function. The average *<Δ>* value of both proteins was found to be < 10 nm (Fig. 4A) and therefore treated as consistent with zero, since globular proteins typically have an intrinsic diameter of a few nanometers.

**Fig. 4.**
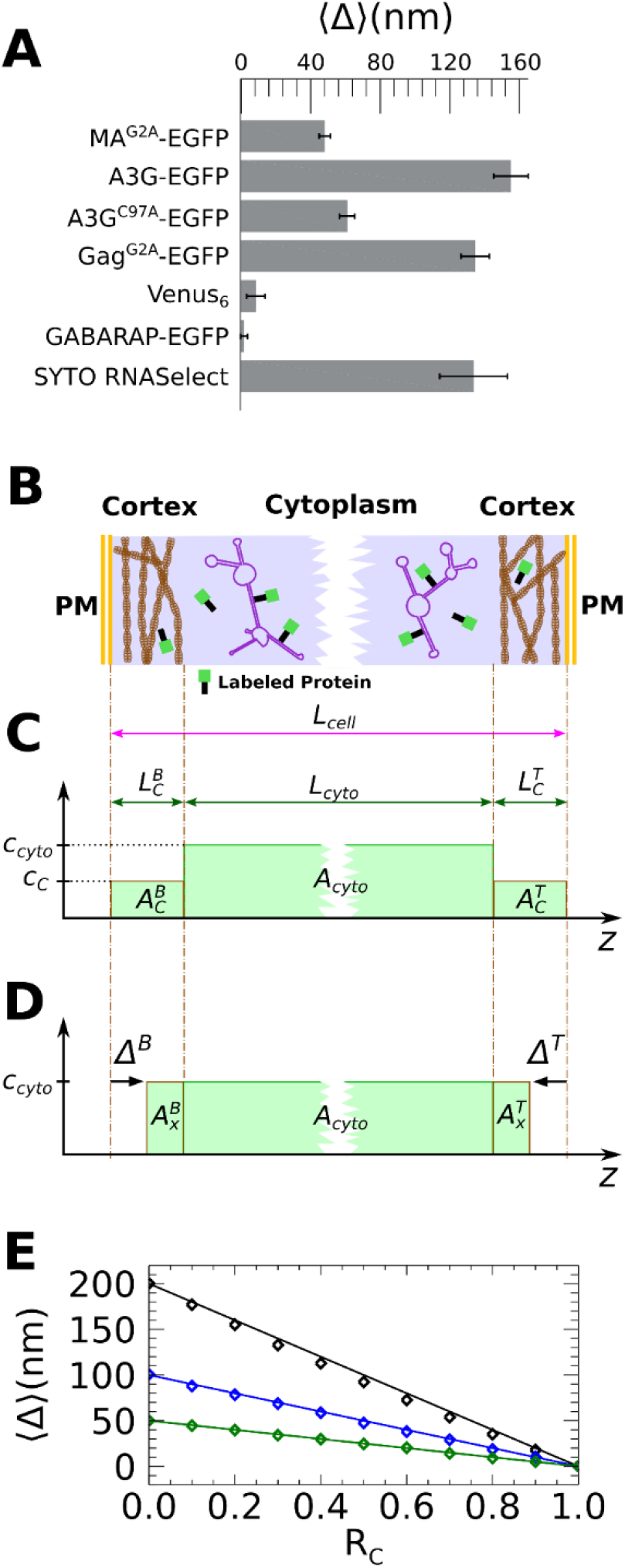
Effective depletion length. A) Average <*Δ*> values in HeLa cells for the indicated proteins and an RNA dye. B) Model of cortical partitioning leading to a SSS^G^ distribution of fluorescence. C) Axial concentration profile corresponding to the model. The concentration drops from *c*_*cyto*_ in the cytoplasm to *c*_*C*_ in the cortex. D) Approximation of the SSS^G^ by a single S^G^ concentration profile. Fitting of the experimental z-scan data to this profile was used to identify apparent depletion lengths *Δ*^B^ and *Δ*^T^. The mCherry concentration profile has been omitted in panels C and D. E) <*Δ*> vs partition coefficient *R*_C_ extracted from simulated DC z-scans for three different cortex lengths (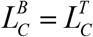 of 200, 100, and 50 nm). The solid lines represent a theoretical model based on Eq. 1.

For comparison, the previously presented depletion lengths from RNA binding proteins (from Fig. 3D) were added to Fig. 4A. Since RNA binding appeared to be important for observing depletion, DC z-scans were performed with the mutant A3G^C97A^-EGFP, which has impaired RNA binding due to mutation of the N-terminal zinc finger (*39*). These measurements revealed a large reduction in *<Δ>* compared to that of wildtype A3G-EGFP (Fig. 4A). Lastly, the *<Δ>* value was measured in mCherry expressing HeLa cells that were stained with the small molecule RNA binding dye, SYTO RNASelect. These measurements revealed that the labeled cellular RNA had a robust depletion length (Fig. 4A).

Based on these results, it was hypothesized that a partial exclusion, or partitioning, of large RNA species from the dense cortical meshwork may be responsible for the observed PM adjacent depletion of RNA binding proteins (Fig. 4B). To determine whether engagement of RNAs at ribosomes might contribute to this effect, DC z-scans were performed in U2OS cells expressing HIV-1 Gag^G2A^-EGFP and mCherry that were treated with puromycin to induce premature chain termination and dislodge ribosomes from mRNAs. Puromycin treatment resulted in *<Δ>* values that were unchanged compared to untreated cells (Fig. S3), suggesting that engagement of mRNAs at ribosomes is not required for cortical partitioning of HIV-1 Gag.

### DC z-scan modeling of cortical partitioning

In a simple model of the cortical partitioning (Fig. 4C), the ratio of the concentration *c*_*C*_ in the cortical layer to the concentration *c*_*cyto*_ in the cytoplasm defines the partition coefficient *R*_*C*_ = *c*_*C*_ / *c*_cyto_. This partitioning of the protein concentration between the cytoplasmic and cortical spaces introduces a SSS^G^ concentration profile for the EGFP-labeled RNA-binding protein (Fig. 4C), which consists of two thin cortical slabs of length 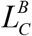 and 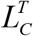 separated by a thick cytoplasmic slab of length *L*_*cyto*_ with a higher protein concentration. As before (Fig. 3), the cortical layer is too narrow to be directly resolved by SC z-scan analysis. However, since the concentration within the cortex layer is reduced (Fig. 4C), the z-scan intensity profile can be effectively approximated by a single slab layer with a width that slightly exceeds the cytoplasmic thickness *L*_*cyto*_ (Fig. 4D). Consequently, the apparent depletion lengths *Δ*^*B*^ and *Δ*^*T*^ will be less than the corresponding cortex thickness 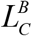 and 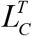 in a manner that depends on *R*_*C*_. In z-scan experiments, this dependence is well approximated by

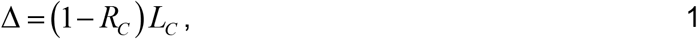

where the superscript has been omitted. Eq. 1 is derived by preserving the integrated fluorescent areas (i.e., *A*_*C*_ = *A*_*x*_) between the partitioning model (Fig. 4C) and the fit model (Fig. 4D). See Supplemental Text 1 for details. This model (Fig. 4E, solid lines) is in excellent agreement with the results from fitting simulated DC z-scan traces of an SSS^G^-S^CH^ geometry to the S^CH^-S^G^ DC z-scan model (Fig. S4). Eq. 1 was used to estimate the partition coefficient of proteins from their measured depletion lengths (Fig. 3D) using the experimentally determined cortex thicknesses *L*_*C*_. *R*_*C*_ = 0.76 ± 0.03, 0.33 ± 0.09, and 0.22 ± 0.1 for MA^G2A^-EGFP, Gag^G2A^- EGFP, and A3G-EGFP, respectively, were determined from HeLa cells. Similar values (0.75 ± 0.04, 0.39 ± 0.08, and 0.13 ± 0.13) were obtained from the measurements in U2OS cells.

### Direct observation of the partition coefficient in the periphery of HeLa cells

Because these estimates of the partition coefficient *R*_*C*_ rely on measurements of *Δ* and *L*_*C*_, which is a new method, a second approach was developed to directly estimate *R*_*C*_ in thin cytoplasmic regions. To illustrate this approach, a ratio image of the red and green detection channels was calculated from a two-photon z-stack acquired in cells co-expressing HIV-1 Gag^G2A^-EGFP and mCherry (Fig. S6). The ratio image reveals that Gag^G2A^-EGFP is more excluded from thin peripheral regions than thick regions of the cytoplasm when compared with the mCherry reference. A similar effect was previously documented using large fluorescent molecules that were microinjected into the cytoplasm (*15, 40*). Based on the partitioning model presented in Fig. 4C, it is anticipated that this relative exclusion seen in peripheral regions is caused by close opposition of the top and bottom cortical layers in thin cytoplasmic sections. As a control, Gag^G2A^-EGFP was replaced with EGFP alone. The resulting ratio image showed an approximately constant intensity ratio throughout the cytoplasm (Fig. S6), as expected given the lack of cortical partitioning for EGFP.

The same relative exclusion of HIV-1 Gag^G2A^-EGFP compared to mCherry is detected by DC z-scans in thick and thin regions of a HeLa cell (Fig. 5C & D). In contrast, DC z-scans in thick and thin regions of a cell co-expressing EGFP and mCherry indicated that the relative concentrations of the EGFP and mCherry species are identical regardless of cytoplasm thickness (Fig. 5A & B). Next, a sequence of z-scans was taken within a single cell along a line that extended from thick to thin cytoplasmic regions. The axially averaged partition coefficient 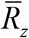, which describes the cumulative partitioning inclusive of both cortical and unpartitioned cytoplasmic layers was constructed from the data (see Supplemental Text 2). Assuming that the cortex thickness *L*_*C*_ and the partition coefficient *R*_C_ are constant throughout the cell as illustrated in Fig. 5E, an analytical expression for 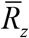 was derived (Eq. S4; see Supplemental Text 2). This theoretical 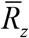 approaches 1 in thick cell sections where the cortical layers are small compared to the cytoplasmic layer, while it approaches *R*_*C*_ in thin cell sections where the unpartitioned layer is small compared to the cortex. While realistic cortices are unlikely to have homogeneous F-actin structures extending from the center to the periphery of cells, (i.e., to have constant values of *R*_*C*_ and *L*_*C*_ in the cell periphery) this model (Fig. 5E) remains useful as baseline against which to assess experimental data.

**Fig. 5:**
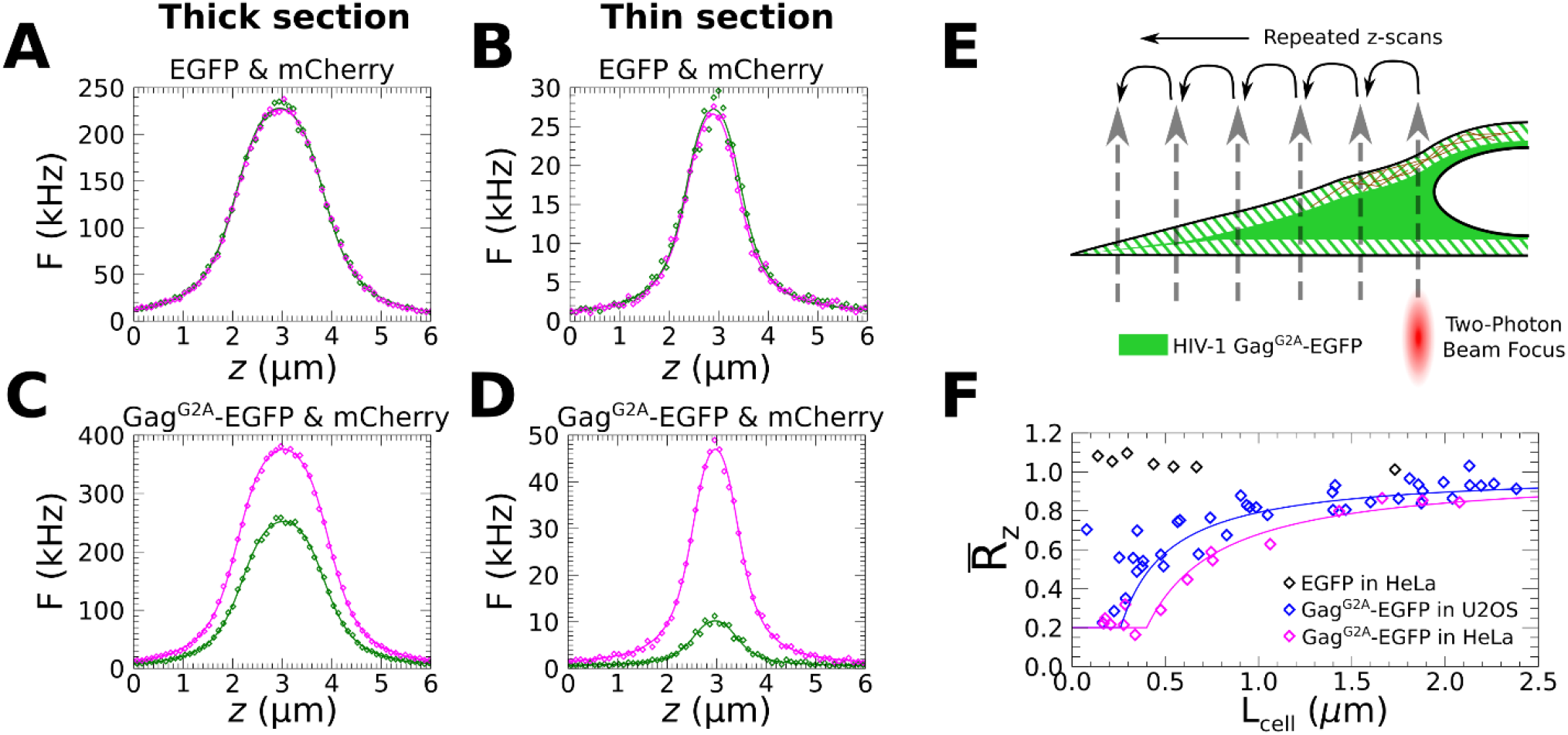
Axially averaged partition coefficient 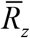 vs cell thickness. A-D) Representative DC z-scan traces in a thin and thick section of a HeLa cell co-expressing EGFP and mCherry (A, B) and representative DC z-scan traces in thin and thick sections of a HeLa cell co-expressing Gag^G2A^-EGFP and mCherry. E) Schematic illustration of DC z-scans taken at multiple cytoplasmic locations varying from thick to thin sections to determine 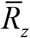 as a function of thickness *L*_*cell*_. F) 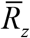 vs *L*_*cell*_. Black diamonds correspond to control measurements in HeLa cells co-expressing EGFP and mCherry. Magenta and blue diamonds correspond to data from HeLa and U2OS cells co-expressing HIV-1 Gag^G2A^-EGFP and mCherry. Magenta and blue lines correspond to Eq. S4 with *L*_*C*_ = 200 nm and *L*_*C*_ = 130 nm, respectively, and *R*_*C*_ = 0.2.

As shown in Fig. 5F, the theoretical 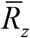 model agrees with the DC z-scan estimates of 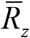 from HeLa cells. The HeLa cell data in Fig. 5F cluster near 0.23 ± 0.02 (SEM, *n* = 6) in thin cell sections. This value is consistent with the *R*_*C*_ value determined above. As a control, the same experiment was carried out in HeLa cells co-expressing EGFP and mCherry. These values remain near 1 independent of thickness (Fig. 5F), as expected due to the lack of cortical partitioning for EGFP. Interestingly, although DC z-scan estimates of 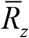 for Gag^G2A^-EGFP in U2OS cells are close to 1 for thick cell sections, they appear to deviate from the model (Eq. S4) in thin cell sections (Fig. 5F). This deviation may reflect the presence of distinct peripheral F-actin structures in this cell line (*28*), which imply changes in the *R*_*C*_ and *L*_*C*_.

### Cortical partitioning and cytoplasmic diffusion are quantitatively related

The cortical partitioning effect documented here supports a model where protein depletion within the cortex is driven by protein association into ribonucleoprotein complexes that are much larger than the typical protein. In such a case, the large *<Δ>* values should be correlated with substantially slowed cytoplasmic diffusion. FCS measurements were performed in HeLa cells in order to estimate the effective diffusion coefficient *D*_*eff*_ of labeled proteins in the cytoplasm (see Materials and Methods and representative FCS curves in Fig. S7). Because FCS data were acquired with the two-photon PSF focused near the middle of thick (>1.5 µm) cytoplasmic sections, the estimates of *D*_*eff*_ correspond to the mobility within the cytoplasm and not the mobility within the actin cortex. In Fig. 6A *<Δ>* values from Fig. 4 were plotted versus the median *Deff* that was obtained from FCS measurements. Two other members of the APOBEC3 protein family, A3A-EGFP and A3C-EGFP, which have differing antiretroviral activities (*41*–*43*), were also measured and added to Fig. 6A. These data show that large *<Δ>* values are associated with remarkably low cytoplasmic mobility. This relationship is summarized by the cartoons in Fig. 6 B&C.

**Fig. 6:**
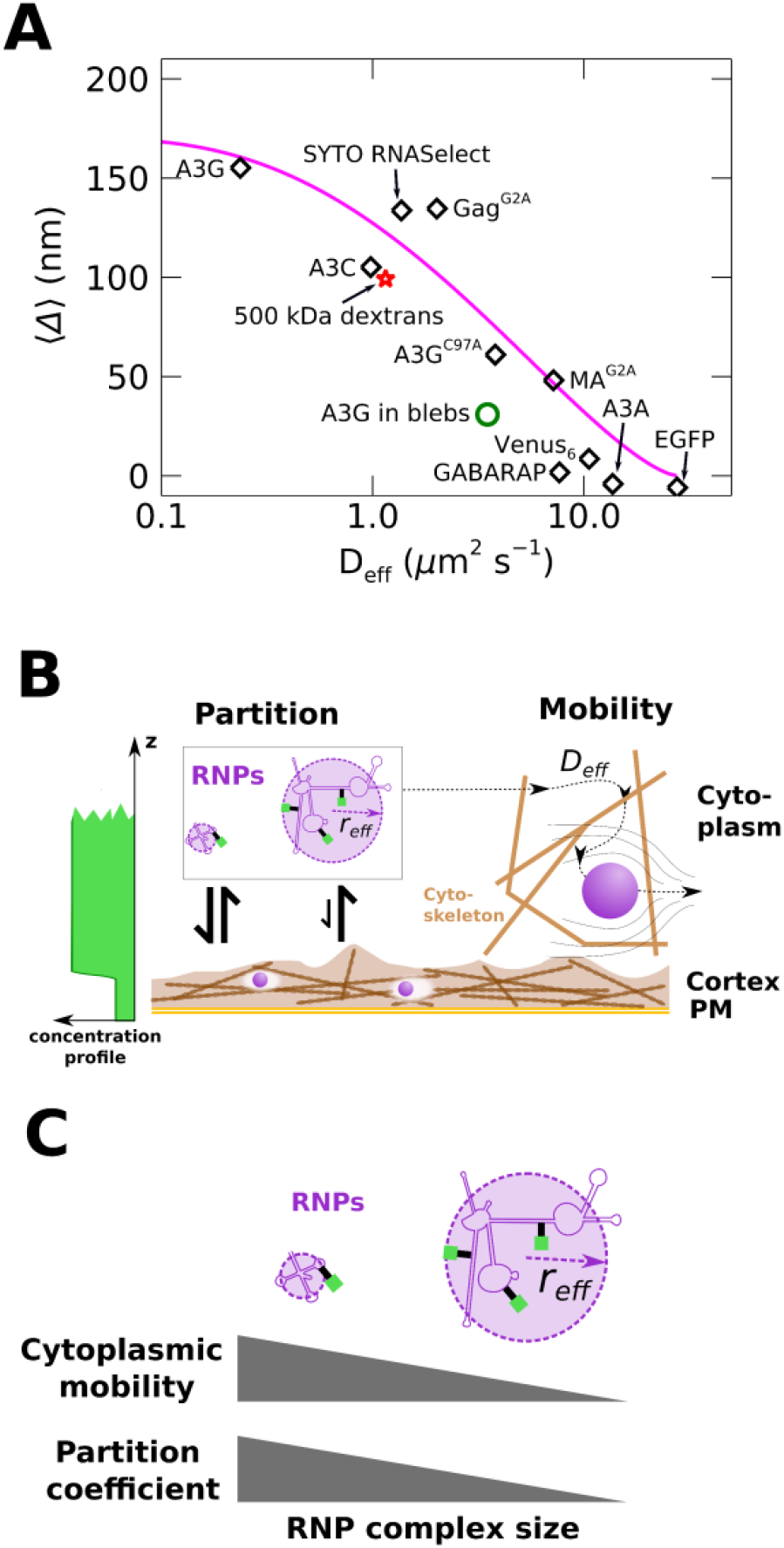
Protein mobility and effective depletion length. A) Depletion length *<Δ>* versus effective diffusion coefficient *D*_*eff*_ in HeLa cells (diamonds). Eq. 4 (magenta curve) is plotted with parameter values as described in the text. A3G-EGFP measured in HeLa cell blebs (green open circle) displayed increased mobility along with decreased effective depletion length. B) Conceptual model of the effects quantified in panel A. The partitioning effect (center left) arises due to hindered entry of large RNP complexes into the cortical mesh, while hindered mobility of large RNP complexes due to macromolecular crowding occurs in the cytoplasm (right). Binding of labeled proteins to partitioned RNP complexes yields the concentration gradient (far left) observed by DC z-scan measurements. C) The cytoplasmic mobility and partition coefficient are connected by the average RNP complex radius, as indicated in the graphical depiction and suggested by Eqs. 3 and 4.

The average pore diameter of the actin cortex has been reported to range between 50 to 200 nm (*29*), depending on cell type. These values are only slightly larger than or overlap with the size of large RNAs (*6, 7*), so that a partial partitioning of RNAs from the interior of the cortex could be driven by excluded volume interactions. Consequently, the cortical partitioning was described as arising from excluded volume interactions of particles that are allowed to enter spherical voids (*44*–*46*),

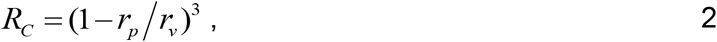

where *r*_*p*_ is the particle radius, and *r*_*v*_ is the void radius. A hindered cytoplasmic diffusion coefficient *D*_*eff*_ was modeled as arising from a size-dependent viscosity imposed by macromolecular crowding within the cytoplasm (*47, 48*) according to

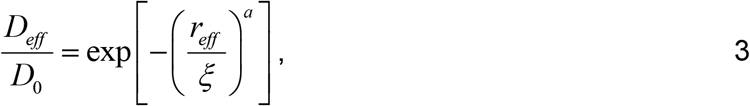

Where *D*_0_ is a reference diffusion coefficient, *r*_*eff*_ is an effective hydrodynamic radius of the diffusing particles, *a* is a scaling exponent of order 1, and *ξ* is a length interpreted as the characteristic distance between points of entanglement within the crowded solution. Using Eqs. 1, 2, & 3, and setting *r*_*eff*_ = *r*_*p*_ yields a relation between *D*_*eff*_ and Δ,

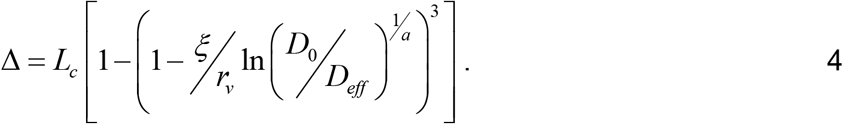

This curve is plotted as the magenta line in Fig. 6A using values *L*_*C*_ = 170 nm, *D*_0_ equal to *D*_*eff*_ measured for EGFP (27.5 µm^2^s^-1^), *r*_*v*_ = 60 nm, *ξ* = 4 nm and *a* = 0.7. The values for *ξ* and *a* are within the range estimated by (*47*) for HeLa cells. *r*_*v*_ is similar to published estimates of the cortical mesh size (*29*). The hydrodynamic radius *r*_*eff*_ of the diffusing protein complexes was calculated (Eq. 3), yielding values in the range of 10 to 40 nm for the slowly diffusing proteins in Fig. 6A (Table S2).

The above model postulates that the cortical partitioning effect arises solely from large hydrodynamic size and is independent of any other interactions between RNPs and the cellular milieu. To check this assumption, DC z-scans and FCS measurements were performed in U2OS cells expressing mCherry that were loaded with 500 kDa green-fluorescent dextrans via electroporation. The measurement of 500 kDa dextrans yielded a relatively large *<Δ>* value and slow diffusion (red star in Fig. 6A), which is consistent with our model. A separate FCS measurement of the 500 kDa dextrans in water estimated a hydrodynamic radius of ∼44 nm, which is comparable to the *r*_*eff*_ values obtained from Eq. 3 for slowly diffusing protein complexes in Fig. 6A (Table S2).

Cytoplasmic crowding in cell associated PM blebs may differ from crowding within undisturbed cytoplasm due at least in part to the lack of a cytoskeletal mesh in the bleb interior. Thus, the mobility of A3G-EGFP in HeLa cell blebs was investigated. FCS measurements (Fig. 6A, open circle) showed an order of magnitude increase in *D*_*eff*_ compared to A3G in the cytoplasm of untreated HeLa cells. The value from blebs is plotted as an open circle in order to emphasize that the model parameters (especially *r*_*v*_ and *ξ*) that describe the rest of the data most likely do not apply to the measurements in blebs.

It was recently reported that the cytoplasmic diffusion coefficient of fluorescent proteins depended on their net charge, with positively charged proteins diffusing slowly. It was suggested that this slow diffusion was due to transient interactions with large, negatively charged cell components such as RNA and the cytoskeleton (*49*). An examination was performed to determine whether the *D*_*eff*_ values in Fig. 6A might be explained by the net charge of each protein-FP fusion. As shown in Fig. S8, the mobility of the proteins in Fig. 6A was only weakly correlated with net charge. Thus, the mobility of these proteins in living cells is likely to be better understood as arising predominantly from their hydrodynamic size as a consequence of their RNA binding function.

## Discussion

In this study, extension of the DC z-scan microscopy technique to measurements of protein localization at sub-diffraction length scales provided the foundation for identifying the cortical partitioning effect and allowed its quantitative description through the effective depletion length *Δ*. Implementation of the DC z-scan measurements using two-photon microscopy eliminates systematic biases, such as chromatic aberration, from the collected data. This absence of biases together with accurate models describing the DC z-scan intensity trace identifies subtle changes in the distribution of fluorescently labeled proteins along the scan path. DC z-scan measurements indicated an apparent restriction in the cytoplasmic volume occupied by RNA binding proteins (e.g., A3G and Gag^G2A^). This depletion effect was markedly reduced in cell blebs and cells treated with latrunculin A, and was observed to be cell-type dependent in a manner that correlated with DC z-scan estimates of cortex thickness in adherent interphase HeLa and U2OS cells. Thus, the depletion effect is interpreted as being a partitioning, or partial exclusion, of these proteins from the volume of the actin cortex. Furthermore, *Δ* values measured for proteins with and without RNA-binding function suggest that the cortical partitioning is driven by association into RNP complexes with large hydrodynamic size and is not associated with protein molecular mass. The independence of *Δ* from protein mass implies that these RNP complexes have hydrodynamic radii far larger than those attained by globular proteins. This conclusion is supported by measurements of the diffusion coefficient, which shows that a marked decrease in mobility is associated with proteins that have the largest values of *Δ* (Fig. 6).

In vitro and in vivo (*50*) observations of actin assemblies suggest that gel-like properties are nearly ubiquitous, including for the actin cortex specifically (*51*). The cortical partitioning effect may be analogous to solute partitioning from the interior of gels, a well-known chemical phenomenon (*52, 53*). Furthermore, the subcellular heterogeneity driven by the dense cortical environment is reminiscent of those molecular crowding effects which have been reported to drive association or phase separations in cells (*54, 55*). This suggests that the crowded environment within gels may drive biophysical phenomena similar to, but distinct from those associated with molecular crowding in the cytoplasm. The observation of cortical partitioning described in this report builds on earlier work that found strong exclusion of labeled colloidal particles with radii exceeding ∼10 nm from distal regions of mammalian cell cytoplasm, and regions near the centrosome (*15, 40*). Subsequent studies documented that the distal excluding regions were associated with a dense cytoskeletal mesh and were thinner than non-excluding regions (*56*). These and other observations (*44*) have suggested that this partitioning was driven by the cytoskeletal meshwork.

The cortical partitioning of RNPs described in the present study also leads to strong exclusion from distal regions of the cytoplasm (Fig. 5 & S6). However, unlike previous studies, which linked partitioning to the actin cytoskeleton in general (*44*), our observations identified the actin cortex as the causative agent of partitioning. Differentiation between the cortical space and the bulk cytoplasm was not previously achieved and probably reflects experimental limitations in these earlier studies, which are overcome by the use of DC z-scan method employed here. Thus, the cortical partitioning of RNPs observed in this work and the distal exclusion of microinjected particles documented previously (*15, 40*) likely are manifestations of the same physical phenomenon, namely the size-dependent partitioning of particles from the actin cortex.

The measurement of the actin cortex thickness, *L*_*C*_, in HeLa (∼200 nm) and U2OS (∼130 nm) cells in our study represents a novel application of the DC z-scan technique. This application was inspired by a conceptually similar approach based on confocal microscopy that was used to estimate a cortex thickness of ∼190 nm in mitotic HeLa cells (*31*) and of 300 – 400 nm in interphase HeLa cells with imposed spherical shapes (*30*). Interestingly, DC z-scan identified a cortical thickness ≤200 nm for adherent interphase HeLa and U2OS. This is consistent with literature reports which indicate that cortical architecture is strongly associated with cell morphology (*28, 30*). In particular, cell adhesion and cell spreading on the glass coverslip can result in a thinning of the actin cortex (*28, 57*). Thus, differing cortical thicknesses reported here and in literature most likely represent inherent differences associated with the disparate cell morphologies studied.

Measurements of the axially averaged partition coefficient 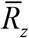 provided further evidence linking partitioning with the cellular cortex by observing 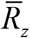 as a function of cell thickness (Fig. 5F). Modeling of 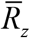 by assuming uniform cortex properties recovered parameters consistent with DC z-scan analysis in the case of HeLa cells. Interestingly, estimates of 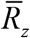 in U2OS cells exhibited more scatter than observed in HeLa cells, particularly in thin cell sections. This may reflect cell-type differences in F-actin assemblies. In EM micrographs, HeLa cells show an unusually uniform cortex, even near the cell periphery, while large structures such as peripheral lamellar actin sheets are more pronounced in other cell types (*28, 58*). Because F-actin structures in lamellae appear distinct from cortical actin fibers, these structures are anticipated to partition large complexes in a manner distinct from the uniform cortex model of 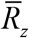.

The strong correlation between the ability of a protein to bind RNA and measured values of <*Δ>* (Fig. 4A) suggests that many RNA-protein complexes are too large to efficiently enter the dense actin cortical meshwork. The cortical partitioning of RNA-binding proteins measured by DC z-scan is thus interpreted as arising from an underlying partitioning associated with the large RNA species with which these proteins associate. In this framework, DC z-scans of RNA interacting proteins indirectly probe the spatial distribution of cellular RNAs. More directly, when cellular RNA was labeled with an RNA-binding fluorescent dye, DC z-scan measurements of <*Δ>* revealed that the RNA itself is partitioned from the cortex with *R*_*C*_ in the range 0.35 – 0.2 (given *L*_*C*_ 170 – 200 nm). This *R*_*C*_ value is interpreted as representing the average *R*_*C*_ of a broad spectrum of differently sized cellular RNAs (i.e., mRNAs, tRNAs, etc) and RNP complexes (e.g., ribosomes) with different degrees of cortical penetration. Similarly, the *<Δ>* values for fluorescently labeled proteins likely represent the average *R*_*C*_ of a heterogenous mix of differently sized RNA complexes to which each protein preferentially binds. Because polysomes can form extremely large structures, the potential for mRNA engagement at ribosomes to drive cortical partitioning was explored by treating cells with puromycin, which causes dissociation of mRNA-ribosome complexes (*59*). This experiment, performed with Gag^G2A^-EGFP, showed no detectable change in *<Δ>* (Fig. S3A), demonstrating that the hydrodynamic size of the probed mRNAs is sufficient for establishing cortical depletion.

FCS measurements were acquired in order to characterize protein mobility in the cytoplasm. The diffusion time of RNA binding proteins revealed that large *<Δ>* values were associated with very low mobility (Fig. 6A), as expected for proteins that are present in large complexes. Slow diffusion of large sized objects in the cytoplasm has been previously reported (*14, 17, 60, 61*), including mRNAs (*62*). Previous studies have suggested that this low cytoplasmic mobility may reflect a hindrance created by cytoskeletal elements (*15, 16, 18, 19, 56, 63, 64*) or by cytoplasmic crowding created due to ribosomes, organelles and proteins (*17, 60, 65*–*67*). Although the precise determinants for cytoplasmic mobility have not been fully characterized, existing literature provides extensive evidence that the cytoplasm cannot be described as a simple fluid (*12, 47, 65, 68*), and it is generally agreed that particles in the range of 10 – 100 nm show a size-dependent drop in cytoplasmic mobility much greater than that predicted by the Stokes-Einstein relation (*14, 17, 18, 60*). In this study, a marked increase in A3G-EGFP mobility in cell-associated blebs (Fig. 6A) was observed. This suggests a role for the cytoskeleton in the mobility of large macromolecular complexes.

To capture the size dependence of RNP diffusion, an empirical hydrodynamic scaling model (Eq. 3) was used. This and similar models have been previously employed to describe diffusion in polymer solutions and in living cells (*47, 48*). This model led to good agreement with our experimental data (Fig. 6A) when combined with a relatively simple description of the size dependence of cortical partitioning (*44*–*46*) (Eq. 2). Although large RNP complexes are the focus of the current work, the physical model should apply to any large, freely diffusing complex. This generality of the cortical partitioning phenomena is supported by data from 500 kDa dextran polymers, which are not native to the cellular milieu but nonetheless show a robust partitioning effect in line with their hydrodynamic size (Fig. 6A). This demonstrates that cortical partitioning may be understood as arising from the simple physical interactions captured by Eqs. 1 – 3 without appeal to complex biochemical interactions that occur in the living cell.

In the cortical partitioning phenomenon, the physical properties of the cortical space appear to drive subcellular segregation of macromolecules in the absence of a clear boundary such as a membrane. Other instances of biomolecule compartmentalization in the absence of membranes have been recognized, such as that observed between P-bodies and the cytoplasm (*69*), and between nucleoli and the nucleoplasm (*70*). In such cases, the distinct physical properties of each subcellular compartment regulate their molecular content (*71*). Partitioning of large complexes from the cortical space is consistent with observations demonstrating that a highly orchestrated coupling between exocytic pathways and cortical actin polymerization is needed to facilitate passage of vesicles through the cortex when the actin fiber density is transiently lowered (*72, 73*). Moreover, the cortical partitioning effect might hinder the replication of enveloped viruses which target large macromolecular complexes to the PM as part of their assembly processes. In the particular case of HIV-1, a large RNA genome must transit the cortical space in order be packaged into the nascent viral particles that assemble at the PM. Recruitment of the viral RNA to assembly sites is driven by the Gag protein, which binds to viral and cellular RNAs (*74*). Because both Gag and the viral RNA are subject to cortical partitioning, the effect may present a barrier to HIV-1 replication that must be overcome during the assembly process.

In conclusion, the extension of DC z-scan microscopy to measure sub-diffraction length scales enabled the observation and characterization of an unexpected cortical partitioning effect that applies to large RNP complexes. A simple quantitative model was introduced to provide a rudimentary foundation for understanding the origins of this effect. The DC z-scan toolkit should be of interest to probe the spatial relation of biomolecules with other cellular structures as well as continue to provide additional insights into the function of the actin cortex.

## Materials and Methods

### Cell Culture and Plasmids

HeLa and U2OS (Numbers CCL-2 and HTB-96, American Type Culture Collection, Manassas, VA) cells were cultured in Dulbecco’s modified Eagle’s medium supplemented with 10% FBS (HyClone Laboratories, Logan, UT) at 37°C and 5% CO_2_. Cells were plated at ∼30% confluency into 8-well chambered slides ∼24 h prior to measurement and transfected with vectors as indicated in the text ∼16 h prior to measurement using GenJet transfection reagent (SignaGen Laboratories, Fredrick, MD) according to the manufacturer’s instructions. The cell growth medium was replaced with phosphate buffered saline immediately prior to measurement. All measurements were performed at room temperature.

HIV-1 Gag (*75*) was ligated into pEGFP-N1 (Clontech, Takara Bio, Mountain View, CA) to generate the HIV-1 Gag-EGFP vector. HIV-1 Gag^G2A^ -EGFP was made using the QuikChange XL Site-Directed Mutagenesis Kit (Stratagene, La Jolla, CA). Venus_6_ was a gift from Steven Vogel (*76*) (Addgene plasmid # 27813; http://n2t.net/addgene:27813; RRID: Addgene_27813). mApple-Lifeact-7 (denoted Lifeact-mApple here) was a gift from Michael Davidson (Addgene plasmid # 54747; http://n2t.net/addgene:54747; RRID: Addgene_54747). All other vectors have been previously described: A3G-EGFP, A3G^C97A^-EGFP, A3C-EGFP, A3A-EGFP (*39*); HIV-1 MA^G2A^-EGFP (*20*); mCherry (*77*); EGFP-HRas (*21*); mCherry-RXR (*78*); GABARAP-EGFP (*79*).

### Instrumentation

DC z-scans and fluorescence fluctuation spectroscopy (FFS) data were acquired on a Zeiss Axiovert 200 microscope modified for 2-photon excitation (*20*). Excitation light was provided by a femtosecond pulsed Ti:S laser (MaiTai or Tsunami models, Spectra Physics, Santa Clara, CA) operated at 1000 nm. This light was focused to a diffraction limited point by a water immersion objective (C-Apochromat, Zeiss, Oberkochen, Germany) with numerical aperture 1.2 and correction collar set according to the glass thickness of the sample holder. Emission light was collected by the same objective and separated from the excitation light by a dichroic filter (740DCSPXR, Chroma Technology, Bellows Falls, VT), then split into red and green detection channels by a low-pass dichroic (FF580-FDi01, Semrock, Rochester, NY). A 515/50nm band pass filter placed before the green channel detector eliminated Fresnel reflected mCherry emission light from the green channel. Emission light in each channel was detected by single photon counting hybrid photomultipliers (HPM-100-40, Becker and Hickl, Berlin, Germany), recorded by hardware (FastFLIM, ISS Inc., Champaign, IL or DPC-230, Becker and Hickl, Berlin, Germany), and saved for later analysis using routines written in IDL 8.6 (Harris Geospatial Solutions, Broomfield, CO).

Axial z-scans utilized a piezo stage (PZ2000, ASI, Eugene OR) driven by a voltage waveform from an arbitrary waveform generator (model 33250A, Agilent Technologies, Santa Clara, CA) to move the sample vertically across the two-photon PSF. All z-scans were acquired by driving the piezo stage with a triangle waveform with a 10 s period and peak-to-peak displacement corresponding to 24 µm. Each period contained two z-scans: the first z-scan passed upward through the cell and the second passed downward through the cell. Each z-scan covered a 24 µm range.

XY scanning of the PSF for acquisition of two-photon images was provided by a galvanometer scan head (Yanus IV, FEI Deutschland GmbH, Planegg, Germany) aligned in the two-photon excitation path. XY positioning was driven by a DAC card (3-axis, ISS Inc., Champaign, IL) controlled by SimFCS software (version 3; G-SOFT Inc., Champaign, IL). All z-scans were acquired within 24 µm of the XY scan field center.

### Z-scan modeling

The single-channel (SC) z-scan fluorescence intensity profile *F* (*z*) is determined by the convolution of the axial concentration profile *c* (*z*) of the fluorescently-tagged protein and the radially integrated point spread function (RIPSF) of the two-photon instrument,

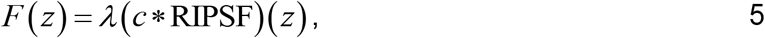

where *λ* is the brightness of the fluorescence tag. The RIPSF was described using the modified Gaussian-Lorentzian model (*22, 33*). This model is specified by a Rayleigh length *z*_*R*_ and exponent *y*, which are calibrated from z-scans through a thin cell section or dye layer (*33*). Calibration values *z*_*R*_ ∼ 0.65 µm and exponent *y* ∼ 1.25 are characteristic of the two-photon instrument used in this study. Evaluation of Eq. 5 for different concentration profiles *c* (*z*) gives rise to an alternative formulation with a fluorescence amplitude *f* and a function v(*z*) describing the shape of the concentration profile (*33*),

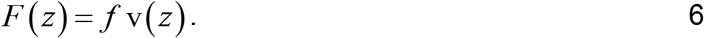

The axially integrated concentration *A* = ⎰ *c* (*z*) *dz* and the axially integrated fluorescence ⋀ = ⎰ *F* (*z*) *dz* are related according to Eq. 5 by

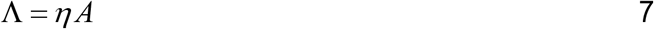

with the constant *η* = *λ* ⎰RIPSF(*z*) *dz*.

### Z-scan and FFS acquisition in live cells

Cells were selected by verifying the desired expression levels for the mCherry and EGFP labeled constructs using 1-photon epi illumination provided by a mercury lamp and using an EGFP/mCherry filter set. Points within cells for z-scan and FFS data collection were chosen where the fluorescence appeared locally (∼3 μm radius) uniform as visualized in epi illumination and where a cell lacked nearby internal structures when visualized by using transmitted light. All measurements of cortical partitioning were performed in the cytoplasm. For data acquisition, the microscope was switched to two-photon mode and the excitation volume was focused to the middle of the cell. At each location chosen, a z-scan acquisition consisted of multiple repeated vertical scans (see Instrumentation section). In most instances, 60 s of z-scan data acquisition (i.e., 12 repeated scan passes) was followed by an optional 60 s of FFS data acquisition at up to 4 unique locations within each cell. Repeated z-scan acquisitions at a given location were measured for a maximum of 10 min in dim samples or where increased statistical precision was desired.

### Effective diffusion coefficient

The effective diffusion coefficient was calculated as (*80*) 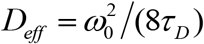, where 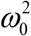 is the 1/ *e*^2^ radial beam waist of the 2-photon excitation light focus and *τ* _*D*_ was estimated by fitting the log-averaged autocorrelation of photon counts *G*(*τ*) to *G*_2*D*_ (*τ*) = *G*_0_ / (1+ (*τ* / *τ*_*D*_))^-1^ + *G*_*offset*_. FCS data quality control was performed by redacting data where *G*_*offset*_ showed significantly negative values or where *τ* _*D*_ indicated by the fit was >50% higher than the time at which the correlation amplitude reached half its initial value.

### Modeling of DC z-scan intensity profiles

The DC fluorescence intensity **F**^**G**^ (*z*) of a z-scan intensity profile of EGFP (G)-labeled protein through a series of horizontal layers, each containing a uniform concentration of labels, is described by (*20*–*22*)

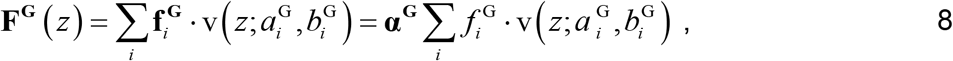

where 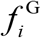 and 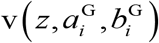 are the fluorescence amplitude and shape function associated with the *i*-th layer. The shape function 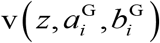 characterizes a uniform slab or S layer starting at 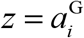 and ending at 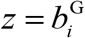, while 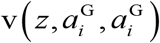 defines an infinitesimally thin or δ layer located at 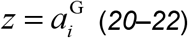. Vector notation is used to describe properties of the green (g) and red (r) channel, i.e. 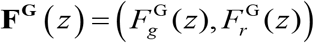 and 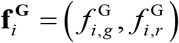. The broad emission spectrum of EGFP is split at the dichroic mirror, resulting in identical shapes of the z-scan profile in both channels. Their intensity ratio is specified by the crosstalk vector **α**^**G**^, which is defined by 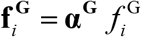, where 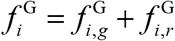 is the sum amplitude of both channels. Similarly, the DC z-scan intensity profiles of the red fluorescent protein mCherry (CH) are given by
0

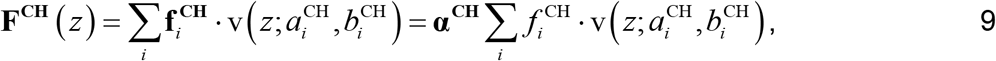

and the presence of both proteins is described by the sum of Eqs. 8 and 9,

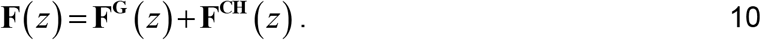

Only a few scan geometries are of interest for modeling of the data in this study. The first uses a single slab layer to describe the distribution of EGFP (X = G) and mCherry (X = CH) labeled species,

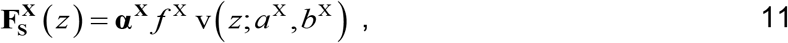

as illustrated in Fig. 1A. The second geometry consists of a delta layer followed by a slab layer and another delta layer, which is referred to as (δSδ) layer model (*20*–*22*). This model was introduced to describe peripheral membrane proteins with the δ layers representing the top and bottom plasma membrane, while the S layer describes the cytoplasmic space (*21*). The DC z-scan (δSδ) model is given by

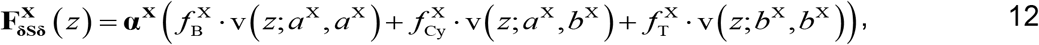

where *a*^X^ and *b*^X^ mark the axial height of the bottom (B) and top (T) membrane (see Fig. 1G). The fluorescence amplitudes of the B, T and cytoplasmic (Cy) layer are given by 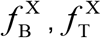, and 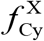.

The DC z-scan of cells co-expressing EGFP and mCherry labeled proteins is described by a superposition of their individual profiles (Eq. 10). The notation X^G^-Y^CH^ was used to describe a model where the z-scan intensity of the EGFP-labeled protein is described by the X profile, while the z-scan intensity of the mCherry-labeled protein is given by the Y profile,

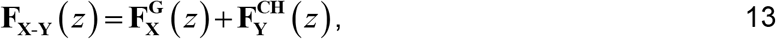

where X and Y denote either the S or δSδ model (Eqs. 11 and 12). For example, the S^G^-S^CH^ model is based on independent slab layers for the EGFP and mCherry-labeled proteins, while the δSδ^G^-S^CH^ model uses the (δSδ) model for the EGFP label and a slab layer for mCherry.

### Production of PM blebs

Bulges along the plasma membrane (i.e., blebs) were induced along the PM of cultured HeLa cells by adding 10-15% ethanol (*25*) or 5 µM cytochalasin D (Cayman Chemical, Ann Arbor, MI) to the cell growth medium and incubating for ∼15 min at 37°C and 5% CO_2_. The drug-treated cells were subsequently analyzed by transmitted light microscopy to assess the number and size of blebs with the goal of producing blebs with ∼5µm diameter in sufficient quantity for measurement. If insufficient blebbing was observed, cells were returned to the incubator for ∼10 min before re-checking. Once sufficient blebbing was obtained, the growth medium with drug was exchanged by PBS containing the same concentration of drug immediately before measurement. For the conditions used to generate ethanol induced blebs, it was observed that a near normal HeLa cell morphology was obtained after ethanol containing media of blebbing cells was exchanged for clean media and cells were incubated for ∼1 h.

Although rapid bleb retraction is often observed in live cell cultures, the blebs produced by ethanol and cytochalasin D treatment were stable during z-scan acquisitions. Bleb stability is likely due to the impairment of the F-actin assemblies that are required for bleb retraction. Over the course of the experiment (several hours), the population of ethanol induced blebs were observed to coalesce into one or several large blebs per cell. Such large blebs were not measured since they tended to contain large internal structures which confound z-scan analysis.

### Puromycin treatment of U2OS cells

Puromycin (Sigma-Aldrich, St. Louis, MO) in water was added to the cell growth medium of U2OS cells to a final concentration of 100 µg/mL. The cells were cultured with the drug for approximately 1 h before the start of measurements. For z-scans, the cell growth medium was exchanged for PBS containing the same concentration of puromycin.

### Latrunculin A (LA) treatment of HeLa cells

LA (Cayman Chemical, Ann Arbor, MI) in DMSO was added to the cell growth medium of HeLa cells to a concentration of 0.15 µg/mL. The cells were cultured with the drug for approximately 1 h before the start of measurements. The experimental LA concentration corresponded to a DMSO concentration of 0.1 %. For z-scans, the cell growth medium was exchanged for PBS containing the same concentration of LA.

### Loading of labeled dextran in U2OS cells

U2OS cells previously transfected with mCherry were trypsinized from a T-25 flask near confluency (∼ 3x10^6^ cells total) and pelleted via low-speed centrifugation. Cells were resuspended in 30 µL PBS. Dextran fluorescein (500 kDa MW, Life Technologies Corp. Eugene, OR) was diluted in PBS to a concentration of 10 mg/mL. A volume of 30 µL of this dextran solution was added to the cells. Next, cells were electroporated using the MammoZapper electroporation system (Tritech Research, Inc. Los Angeles, CA) according to the manufacturer’s recommended protocol. To remove excess dextran, 1 mL growth medium was added to the cells, which was followed by pelleting at low-speed centrifugation. This wash step was repeated a total of three times. Cells were then plated into 8-well chambered slides and incubated for 24 h. The cell growth medium was exchanged with PBS immediately prior to measurement.

### Imaging of TetraSpeck beads

0.1 µm TetraSpeck beads (Thermo Fisher Scientific, Waltham, MA) were diluted in water and immobilized on imaging slides by addition of PBS buffer followed by quick washing with clean buffer. For measurement, an isolated bead was selected and a z-stack of 200 XY images was acquired with z-steps of 50 nm. Each XY image (256x256 pixels) was taken with a pixel resolution of 26.6 nm.

### SYTO RNASelect cell stain

Live cells were stained with SYTO RNASelect (Thermo Fisher Scientific, Waltham, MA) by diluting the stock dye 1000x (5 µM final concentration) into the cell culture medium followed by incubation for 1 h at 37 °C and 5% CO_2_. For measurement, the growth medium with SYTO RNASelect was replaced by PBS containing the same concentration of the dye.

### Z-scan redaction and statistical analysis

Despite the selection of scan locations as described earlier, some of the measured z-scan intensity profiles were not adequately described by our fit model. This was attributed to scans that inadvertently passed through complex intracellular structures. Since these structures are not captured by the model framework, the spatial autocorrelation of the normalized residual from the fit was calculated to identify patterns that are indicative of the presence of extraneous cellular features not described by the fit model. Scans with a correlation amplitude exceeding a threshold value were removed. Furthermore, z-scan measurements of the cortex thickness were selected to have a cell thickness *L*_*cell*_ ≥ 2.5 µm and a membrane fraction (*22*) *m* ≥ 0.5 in order to ensure the absence of biases in the fitted values as supported by our study using simulated DC z-scans (Fig. S4). Quality control for the z-scan fits of cortex thickness was provided by averaging the magnitude of the normalized residuals within ±1 µm of the fitted PM locations. Scans where these averaged residuals exceeded 2.5 in either the red or green channel were redacted.

A typical 60 s z-scan acquisition at a single location contained 12 repeated scans, which were fit individually, yielding 12 estimates of each parameter of interest. This set of parameters served to calculate the average effective depletion length Δ_*i*_ and its standard deviation *σ* _*i*_ at the *i*-th z-scan location. An average over multiple measurement locations, denoted by angle brackets, was estimated by the weighted average ⟨ Δ ⟩ over all locations with 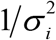 as weight. The uncertainty in the weighted average was computed using the dispersion corrected standard error of the mean (SEM),

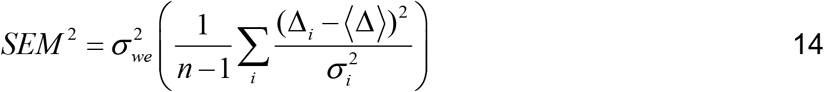

where 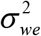 is the variance of the weighted average. Since it was observed that ⟨ Δ ⟩ at the top (T) and bottom (B) PM interfaces are in agreement (⟨ Δ^*T*^⟩ ≈ ⟨ Δ^*B*^⟩), data from both interfaces were grouped together when calculating ⟨ Δ ⟩. This is indicated by dropping the *T* and *B* superscript. Note that when datasets are grouped in this manner, estimates of *Δ*^*T*^ and *Δ*^*B*^ are considered as independent, which is reflected by a doubling of the *n* value when calculating SEM.

### Simulations of z-scan traces

Simulated DC z-scan traces were obtained following a previously described procedure (*20*) to evaluate models and estimate the statistical uncertainty in DC z-scan fit results. Simulated DC z-scan traces were fit using the same algorithms that were applied to experimental data. All data derived from simulations represent the average result from fits of *n* = 1000 z-scan traces.

## Non-author contributions

The authors thank Yan Chen for help with logistics and experiment planning.

## Author contributions

CIA, JDM, and SRK developed the dual color z-scan method. CIA and SRK performed experiments and acquired the data. LMM helped interpret the data and guide the project. CIA and JDM wrote analysis code, performed data analysis, and drafted the manuscript. All authors reviewed and revised the manuscript.

## Data and materials availability

Data and z-scan analysis code are available at Dryad: https://doi.org/10.1016/j.bpj.2019.12.002

