## Supplementary Information for "Partitioning of ribonucleoprotein complexes from the cellular actin cortex"

Science Advances

AAAS

**This PDF file includes:**

Supplemental Text1 & 2

Figures S1 to S8

Tables S1 & S2

### Supplemental Text 1: Derivation of Eq 1

Eq. 1 is derived by preserving the integrated area Λ under the z-scan intensity profile between the SSS$^G$ model and the fitted S$^G$ model. Since Λ and the axially integrated concentration A are directly proportional (Eq. **Error! Reference source not found.**), the concentration areas of both models (Fig. 4C and D) must match. In particular, the cortical concentration area $A_C^i = L_C^i \cdot c_C$ (Fig. 4C) has to be equal to the added extra area $A_x^i = \left(L_C^i - \Delta^i\right) c_{cyto}$ (Fig. 4D) at both the top (i = T) and bottom (i = B) of the cell. Thus, using $R_C = c_C / c_{\text{cyto}}$ we get $\left(L_C^i - \Delta^i\right) c_{\text{cyto}} = L_C^i R_C c_{\text{cyto}}$ , which is equivalent to Eq.1.

**Supplemental Text 2: The axially averaged partition coefficient $\bar{R}_z$ versus cell thickness**

This section first derives a relation between $\bar{R}_z$ and the cell thickness $L_{cell}$, which was used to model the data of Fig. 4F. Subsequently, the experimental determination of $\bar{R}_z$ and $L_{cell}$ from DC z-scan data is described.

The partial exclusion of fluorescently labeled protein from the cortex reduces the fluorescence area $\Lambda^G$ as compared to the area $\Lambda^G_{ref}$ corresponding to the hypothetical case of no exclusion. The ratio of these two areas is denoted as the axially averaged partition coefficient,

$$\bar{R}_z = \frac{\Lambda^G}{\Lambda^G_{\mathrm{ref}}} = \frac{A^G}{A^G_{\mathrm{ref}}} = \frac{L_{cyto} c_{cyto} + 2 L_C R_C c_{cyto}}{L_{cell} c_{cyto}}, \qquad \text{S1}$$

where the proportionality between $\Lambda$ and $A$ (Eq. 7) was used to express the ratio in terms of the integrated concentration areas. Since the cell thickness is defined by (Fig. 3B),

$$L_{cell} = L_{cyto} + L_C^T + L_C^B = L_{cyto} + 2L_C, \qquad \text{S2}$$

where $L_C = (L_C^T + L_C^B)/2$ defines the average cortex thickness, the axially averaged partition coefficient simplifies to

$$\bar{R}_z = 1 + (R_C - 1)\frac{2L_C}{L_{cell}} \qquad \text{S3}$$

This equation predicts $\bar{R}_z \to 1$ for very thick sections ($L_{cyto} \gg L_C$). As the cell thickness $L_{cell}$ decreases, the value of $\bar{R}_z$ drops. It reaches a limiting value of $\bar{R}_z = R_C$ for very thin cell section composed entirely of cortex $L_{cell} = 2L_C$. Thus, DC z-scans in thin regions near the cell perimeter combined with DC z-scans in thick regions of the same cells allow direct estimation of the partition coefficient.

While this relatively simple model sets a minimum cell thickness of $L_{cell} = 2L_C$, where $L_C$ denotes the average cortical thickness in a given cell type, in practice very thin cellular regions were found where the realized cell thickness is less than this value, $L_{cell} < 2L_C$. This implies that in these regions of the cell the local cortex thickness is less than $L_C$. Assuming that these thin sections are composed entirely of cortex, Eq. S3 is generalized to,

$$\bar{R}_z = \begin{cases} R_C & \text{if} \quad L_{cell} < 2L_C \\ 1 + (R_C - 1)\dfrac{2L_C}{L_{cell}} & \text{if} \quad L_{cell} > 2L_C \end{cases} . \qquad \text{S4}$$

Experimental determination of $\bar{R}_z$ from DC z-scans in thin and thick regions of the cell is reflected in the decrease of the integrated fluorescence area $\Lambda^G$ as a function of cell thickness (Eq. S1). To quantify the effect and utilize Eq. S1, an estimate was needed for the reference area $\Lambda_{ref}^G = L_{cell} g_{cyto}^G$, where $g_{cyto}^G = c_{cyto}^G \eta$ (Eqs. 7 and S1). The cell thickness $L_{cell}$ was determined from the mCherry signal in the red detection channel as described in the next paragraph, while the amplitude $g_{cyto}^G$ was estimated from a scan through a thick cell section ($L_{cell}$ > 1.5 µm) where $g_{cyto}^G$ can be reliably extracted from fitting the DC z-scan traces.

Determination of the thickness $L_{cell}$ from DC z-scan data of a cell section that is thick with respect to the RIPSF width is obtained by a direct fit of the intensity profile from the mCherry signal to a slab model (Eq. 11) determines the cell thickness by $L_{cell}$ =

$b^{CH} - a^{CH}$. For cell sections that are thin with respect to the RIPSF, cell thickness cannot be reliably determined from fitting to a slab model (*32*). However, because the fluorescent protein mCherry is distributed uniformly throughout the cell interior, its z-scan intensity provides a measure of local cell thickness $L_{\text{cell}}$. In this manner the cell thickness in thin sections can be determined from the integrated fluorescence $\Lambda^{CH}$ of the intensity profile. $\Lambda^{CH}$ is proportional to the integrated concentration area $A^{CH}$ (Eq. 7), which is $L_{cell} c^{CH}$ for the slab geometry, where the mCherry concentration $c^{CH}$ is assumed constant throughout the cell. Thus, the ratio of the integrated intensity profiles from a thick and a thin section measured in the same cell equals their thickness ratio, $\Lambda_{thin}^{CH} / \Lambda_{thick}^{CH} = L_{cell,thin} / L_{cell,thick}$, which was used to determine the thickness of thin cell sections. Although previous work (*32*) defined the cutoff between thick and thin cell lections as $L_{\text{cell}}$ = 0.5 μm, a more conservative value of 1.5 μm was used in this study to improve the precision of length measurements.

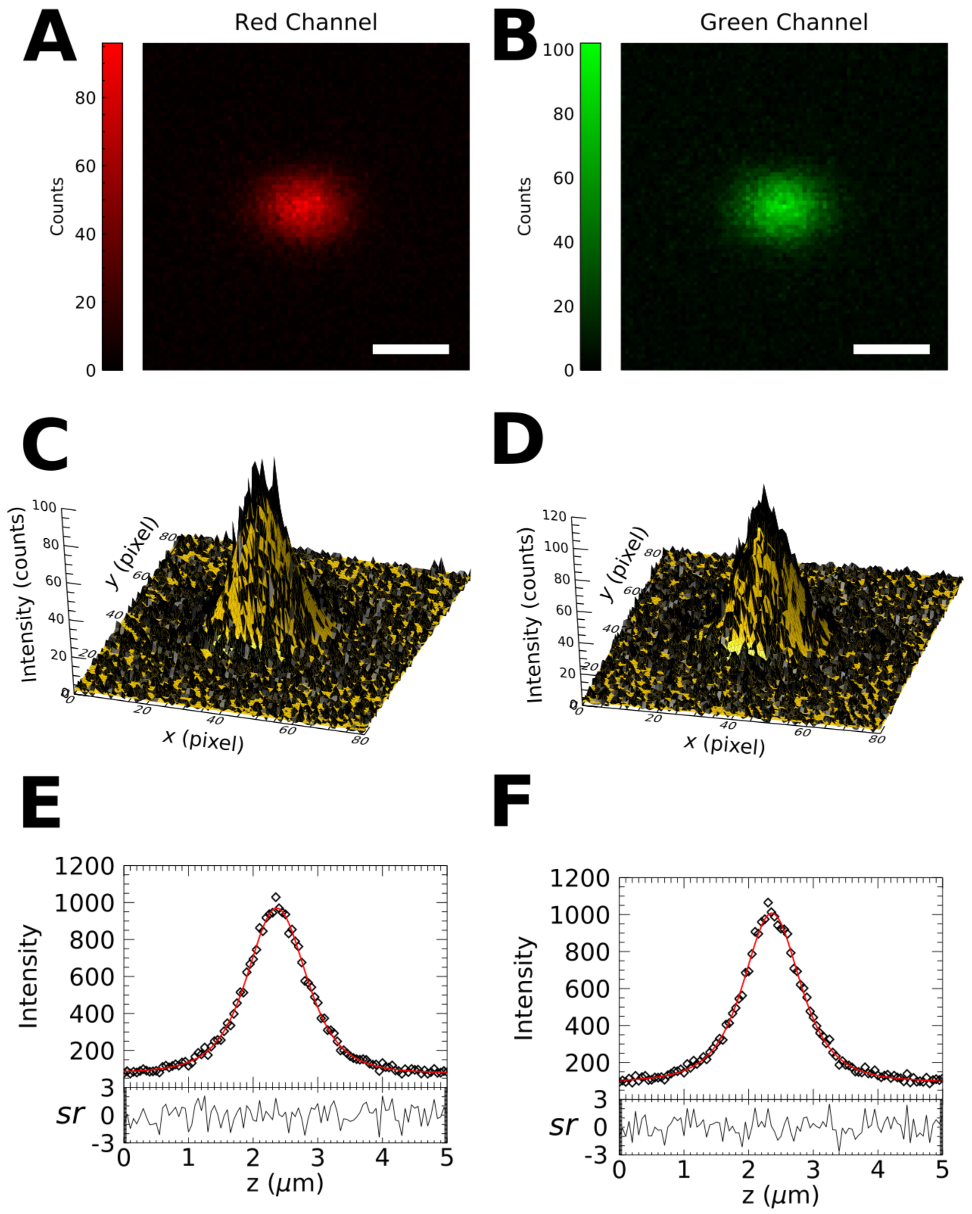


**Figure S1: Absence of chromatic aberrations in the two-photon PSF.** A z-stack of TetraSpeck beads was acquired on the same microscope used for DC z-scan data collection. (A) & (B) Images of a single 0.1 μm TetraSpeck bead in the red and green channels, respectively. Scale bar = 500 nm. Images represent the data after summing over the Z-dimension. (C) & (D) The intensity profiles from panels A & B are fit to 2D Gaussian functions to localize the bead in the XY-dimensions in both channels. The red and green localizations in X and Y differ by -1.5 ± 2.3 nm and 1.4 ± 1.7 nm, respectively. (E) & (F) The z-stack is summed over the XY dimensions to produce a radially integrated z-profile of the bead in the red and green channels, respectively (diamonds). These z-scan profiles are fit to δ layer models (red solid curve) to localize the bead in the vertical dimension in both channels. The red and green vertical localizations of the bead differ by 6.5 ± 6.5 nm.

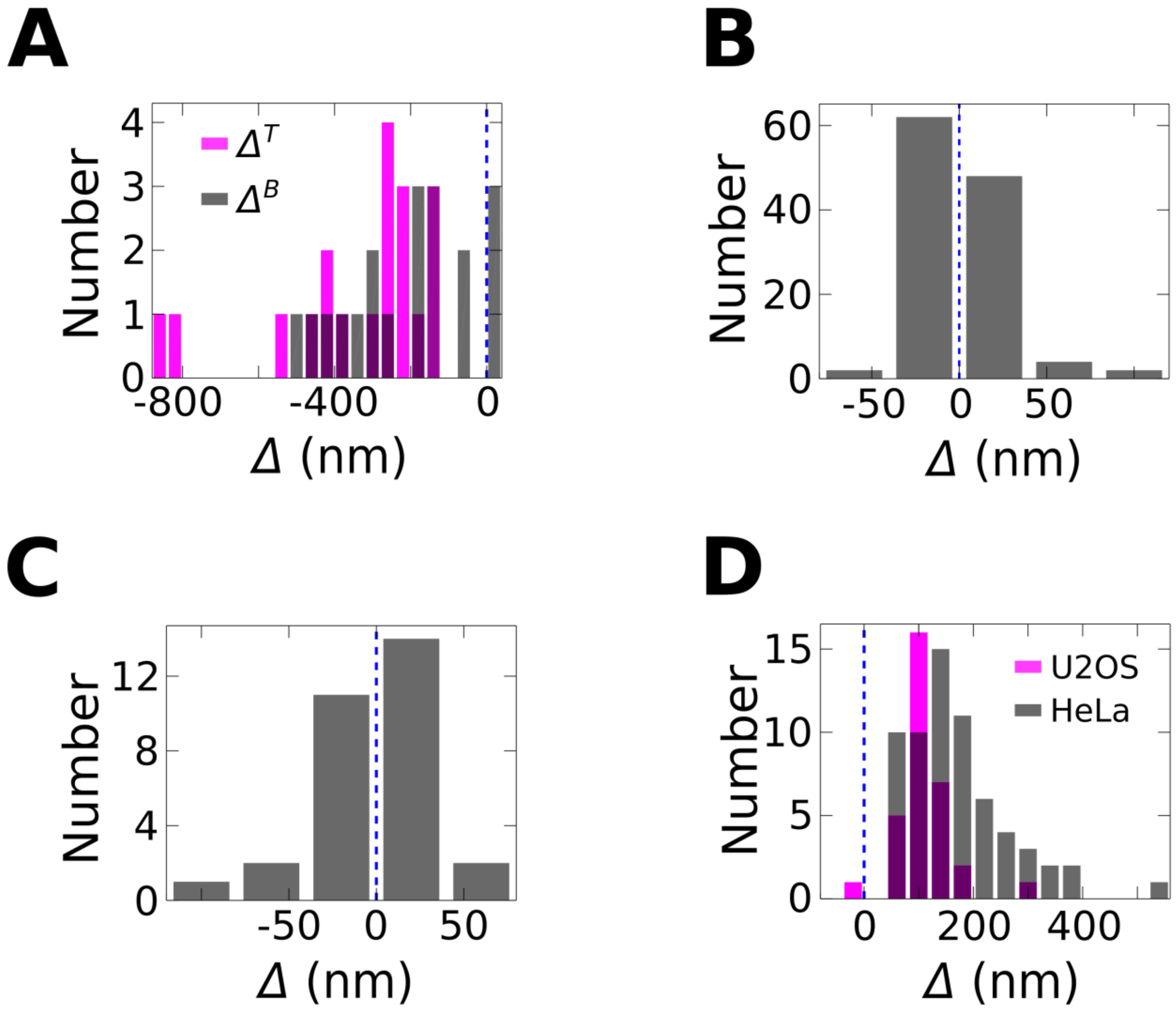


**Fig. S2: Scaled Histograms of *Δ*.** The histograms from Fig. 1 are reproduced with independently scaled horizontal axes. A) EGFP and mCherry-RXR, B) EGFP and mCherry, C) EGFP-HRas and mCherry, and D) A3G-EGFP and mCherry.

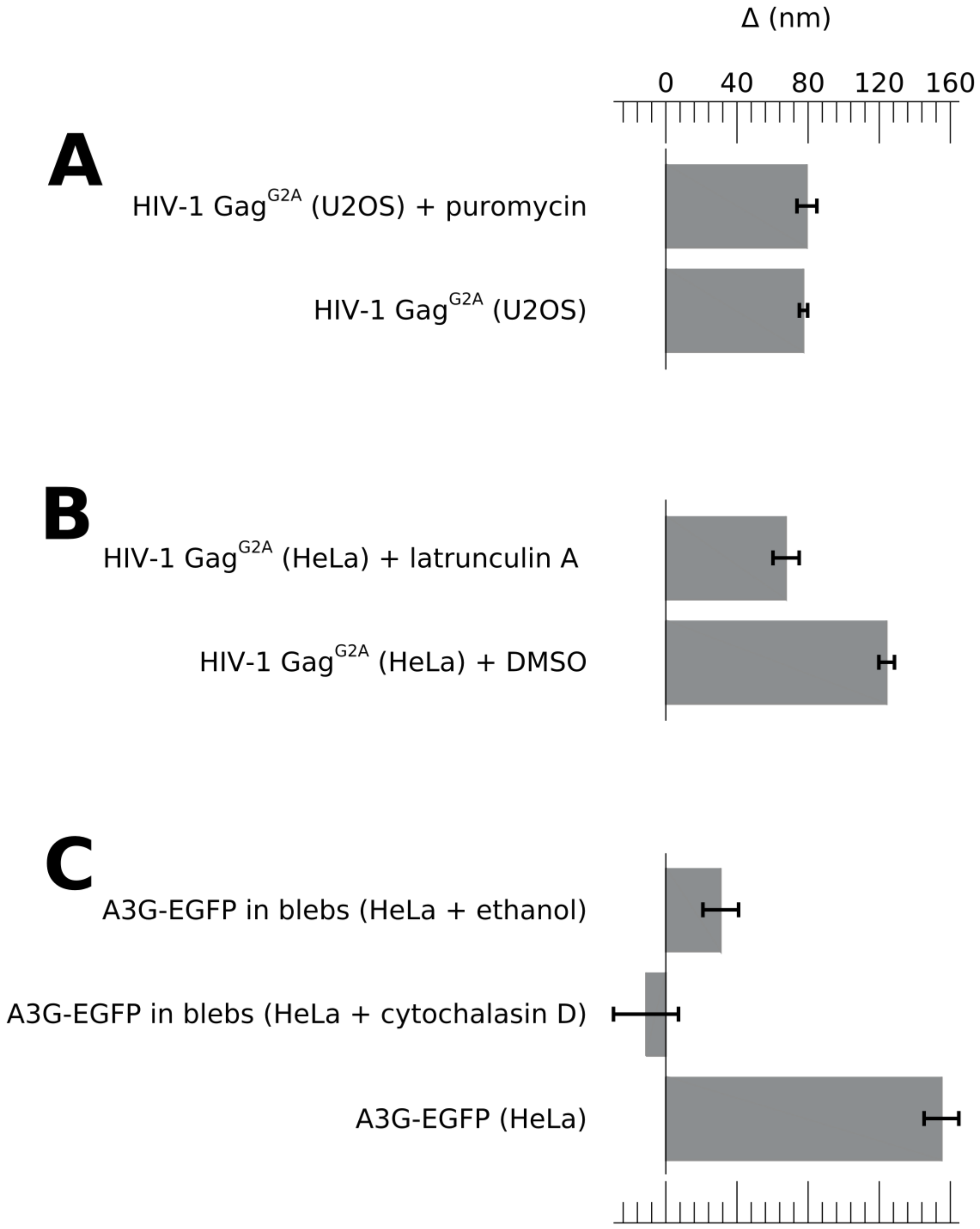


**Figure S3: Depletion length in cells treated with puromycin and F-actin disrupting agents.** A) DC z-scan measurements of HIV-1 $Gag^{G2A}$ in the cytoplasm of U2OS cells treated with 100 µg/mL puromycin yielded the same *Δ* values as measurements performed in untreated cells. B) DC z-scan measurements in the cytoplasm of HeLa cells treated with 0.15 µg/mL latrunculin A showed significantly reduced *Δ* values compared to a solvent control. C) DC z-scan measurements of A3G-EGFP in cell associated blebs generated by treating HeLa cells with ethanol or cytochalasin D yielded much lower *Δ* values compared to A3G in the cytoplasm of untreated HeLa cells.

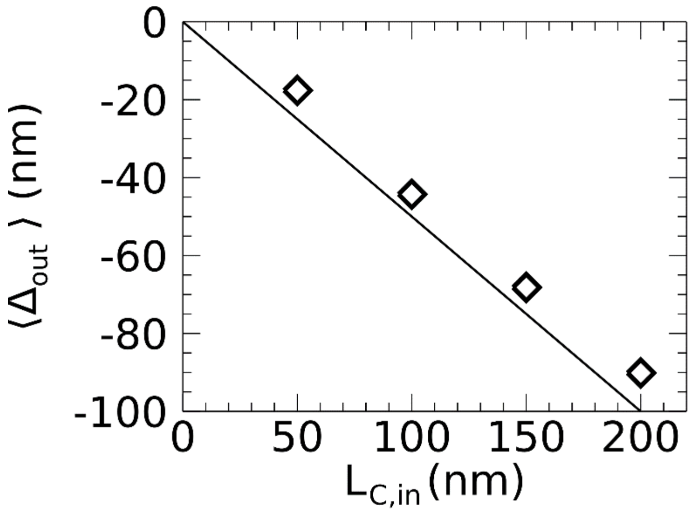


**Fig. S4: Modeling of effective depletion length:** The relation $|\Delta| = L_C / 2$ was tested with simulated traces (see Materials and Methods) of a $SSS^{CH}$- $\delta S\delta^{G}$ model, which were fit to a $\delta S\delta^{CH}$- $\delta S\delta^{G}$ model to extract the effective depletion length $<\Delta_{out}>$. This fitted length is plotted versus the cortex length $L_{C,in}$ chosen for the simulated $SSS^{CH}$ model. Each data point represents the averaged $\Delta_{out}$ from simulations conducted over a range of fluorescent amplitude and $L_{cell}$ parameter space that is similar to the range covered by DC z-scans measured in cells expressing EGFP-HRas and Lifeact-mApple. Additionally, $L_{cell} \geq 2.5$ µm and a membrane fraction $m \geq 0.5$ was chosen to minimize biases in the fitter estimates (20). The line represents the theoretical model, $\Delta_{out} = -L_C / 2$, which agrees with the simulated values to within 10 nm. This result demonstrates that the relation $|\Delta| = L_C / 2$ is experimentally accurate and produces negligible error.

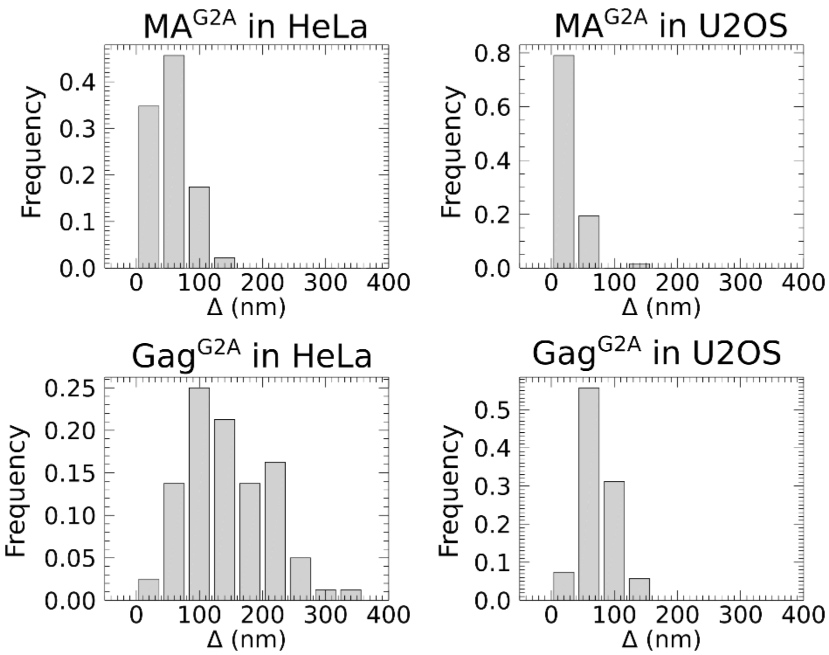


**Fig. S5: Cell-line dependent depletion length.** Histogram of effective depletion lengths measured for the indicated constructs in HeLa and U2OS cells.

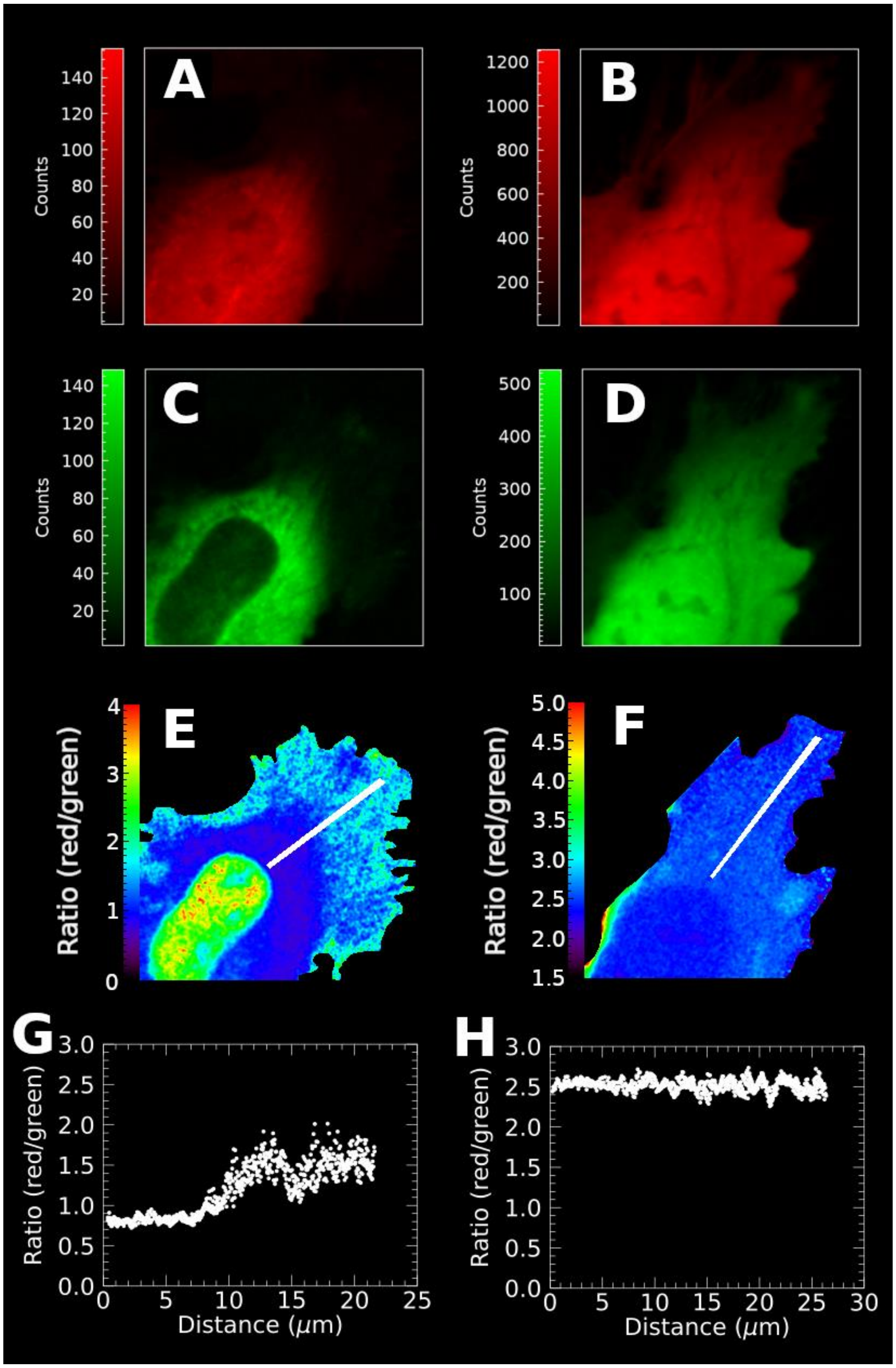


**Figure S6: Two-photon ratio imaging qualitatively detects cortical partitioning in peripheral cytoplasm.** The summed z-stack of the red and green detection channel of a HeLa cell co-expressing Gag$^{G2A}$-EGFP with mCherry (A, C) and EGFP with mCherry (B, D). The red channel (A, B), green channel (C, D), and red to green channel ratio (E, F) images are shown. Line plots of the red/green ratio along the white line (E, F) are shown for the Gag$^{G2A}$-EGFP (G) and EGFP (H) cell. Zero distance represents the point within each stripe that is closest to the cell nucleus. The red/green ratio increases near the periphery of the cell expressing Gag$^{G2A}$-EGFP, indicating the relative exclusion of Gag$^{G2A}$-EGFP from these regions. In contrast, the red/green ratio in the cell expressing EGFP is approximately constant.

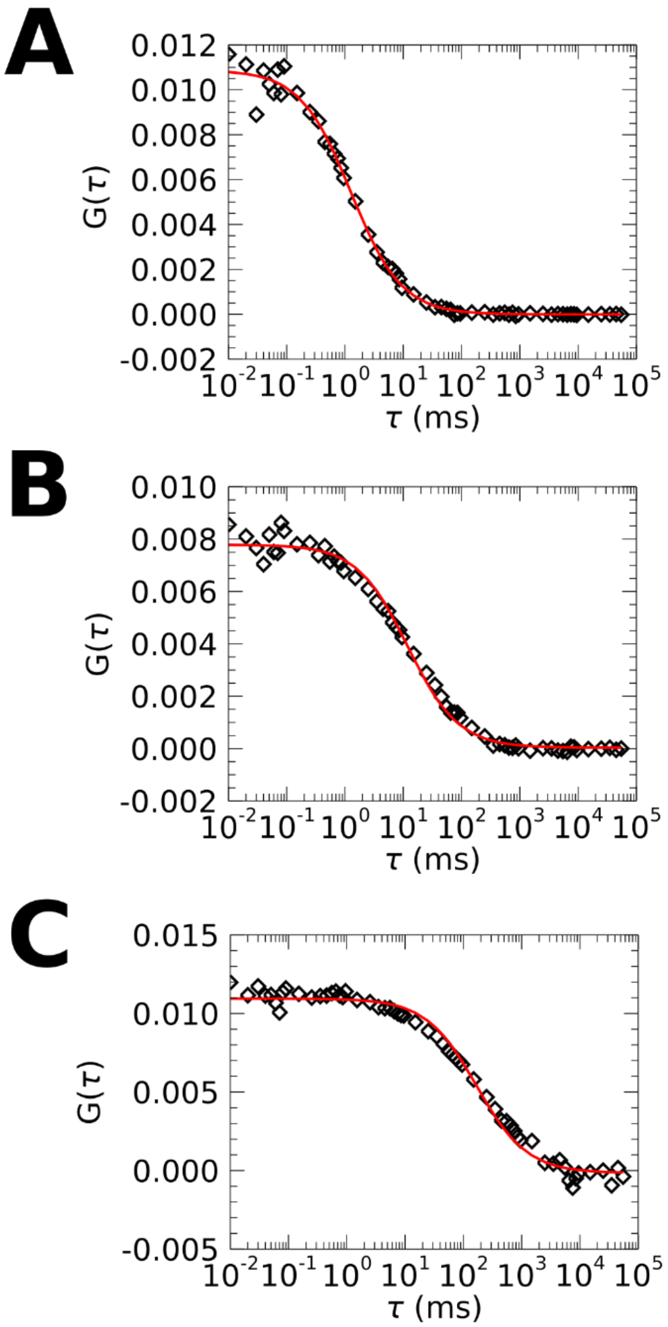


**Figure S7: Representative autocorrelation functions.** Representative autocorrelation functions (ACFs) measured for (A) EGFP, (B) HIV-1 Gag$^{G2A}$-EGFP, and (C) A3G-EGFP in HeLa cells are plotted as black diamonds. A fit to a 2D Gaussian diffusion model is plotted as the red curves. Note that both HIV-1 Gag$^{G2A}$ -EGFP and A3G-EGFP show a slightly broadened decay of the autocorrelation curve, which may indicate heterogeneity in the diffusion coefficients within the sample, anomalous diffusion, or both *(81)*. The characteristic diffusion time $\tau_D$ corresponds to the time where the ACF reached half its initial amplitude, which is adequately determined by the simple diffusion model.

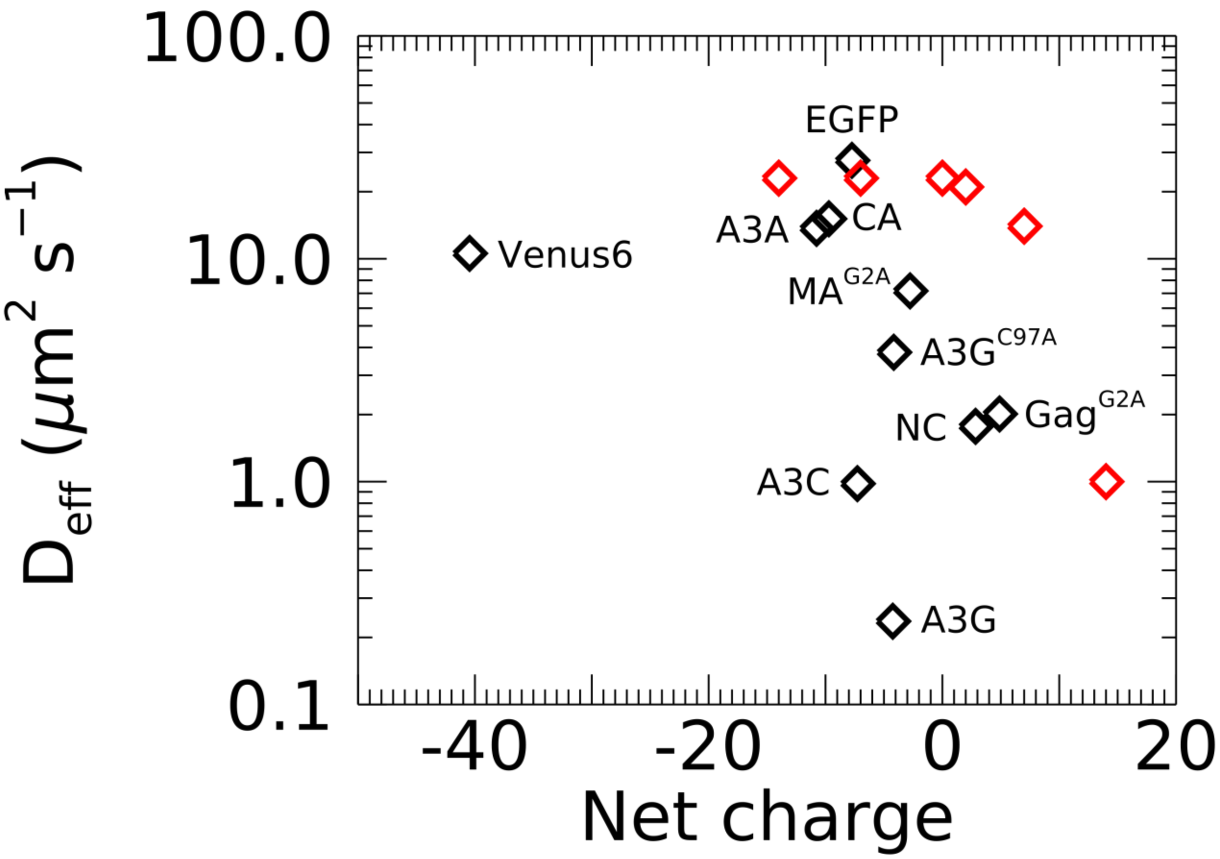


**Fig. S8: Effective diffusion coefficient versus protein net charge (black diamonds).** The red diamonds correspond to diffusion coefficients estimated in *(49)* for fast-diffusing regions of cytoplasm. Net charges were calculated from the amino acid sequence of each protein using an average residue charge at pH 7.4 as described in *(54)*.

| Sample name | STDEV across all z-scans (nm) | average STDEV within single cells (nm) |
|---|---|---|
| EGFP | 20 | 13 |
| HIV-1 Gag$^{G2A}$-EGFP | 66 | 54 |
| A3G-EGFP | 93 | 71 |
| A3G$^{C97A}$-EGFP | 39 | 32 |
| A3A-EGFP | 15 | 12 |
| A3C-EGFP | 44 | 37 |
| SYTO RNASelect | 141 | 73 |
| MA$^{G2A}$ | 28 | 17 |
| Venus$_6$ | 36 | 20 |
| GABARAP-EGFP | 25 | 19 |

**Table S1: Variability in measured depletion values.** The standard deviation of the depletion length $\Delta$ calculated across all z-scans in each dataset is compared to the average standard deviation of the depletion length at distinct locations within the same cell. All values were obtained from HeLa cells. The majority of the variability is accounted for by intracellular heterogeneity.

| | $r_{eff}$ (nm) |
|---|---|
| **Gag$^{G2A}$** | 15.8 |
| **A3G** | 37.1 |
| **A3G$^{C97A}$** | 10.6 |
| **A3A** | 2.4 |
| **A3C** | 22.4 |
| **SytoRNASelect** | 19.2 |
| **NC** | 16.9 |
| **CA** | 1.9 |
| **MA$^{G2A}$** | 6.1 |
| **Venus$_6$** | 3.8 |
| **GABARAP** | 5.7 |

**Table S2: Effective hydrodynamic radii derived from the hydrodynamic scaling model in HeLa cells.** Eq. 3 was solved for $r_{\mathrm{eff}}$ and then evaluated with the model parameter values used in the text and the effective diffusion coefficient $D_{\mathrm{eff}}$ measured for each protein by FCS.
